# Glutamine-regulated VCP acetylation creates a proteostasis vulnerability in RB1-deficient tumor

**DOI:** 10.64898/2026.09.14.751361

**Authors:** Binghui Yang, Ying Cao, Yining Tao, Haoyu Wang, Haoru Dong, Xiyu Yang, Xin He, Bowen Zhao, Kaiyuan Liu, Zongyi Wang, Jiakang Shen, Yan Yan, Qi Zhang, Zhuoying Wang, Yu Lv, Liu Yang, Jing Xu, Tao Zhang, Yingqi Hua, Zhengdong Cai, Haoran Mu, Dongqing Zuo, Hongsheng Wang, Wei Sun

## Abstract

RB1 deficiency defines an aggressive tumor state with limited therapeutic options. Osteosarcoma represents a clinically relevant model of RB1-deficient malignancy, as RB1 alterations are among the most frequent genomic events in this tumor type. However, the non–cell cycle vulnerabilities created by RB1 loss remain incompletely understood. We therefore used osteosarcoma to identify therapeutic dependencies associated with RB1 deficiency. Through high-throughput compound screening, we identified a selective vulnerability of RB1-deficient tumor cells to VCP inhibition. NMS-873 and other VCP-targeting compounds preferentially suppressed RB1-deficient cells across two-dimensional cultures, three-dimensional spheroids, patient-derived organoids, and in vivo tumor models. Mechanistically, VCP inhibition exacerbated proteostasis stress and activated endoplasmic reticulum stress responses in RB1-deficient cells. We further found that VCP function was regulated by glutamine-dependent acetylation, with lysine 615 serving as a dominant acetylation site that modulated sensitivity to VCP inhibition. Glutamine restriction phenocopied, whereas glutamine supplementation partially rescued, the effects of NMS-873 on VCP acetylation and tumor cell growth. Clinical and transcriptomic analyses further linked VCP expression and protein-folding stress programs to aggressive disease features. Together, these findings identify glutamine-regulated VCP acetylation as a metabolic–proteostasis dependency in RB1-deficient tumors and nominate VCP inhibition as a therapeutic strategy for this difficult-to-target tumor state.

**STATEMENT OF SIGNIFICANCE:** Glutamine-regulated VCP acetylation exposes a proteostasis vulnerability in RB1-deficient tumors, supporting VCP inhibition as a therapeutic strategy.

## INTRODUCTION

RB1 deficiency represents a recurrent tumor-suppressor lesion that defines an aggressive and difficult-to-target tumor state. Although RB1 loss has classically been linked to deregulated cell-cycle progression, emerging evidence suggests that it may also create additional dependencies that extend beyond canonical cell-cycle control. Osteosarcoma provides a clinically relevant model for investigating such vulnerabilities, as RB1 alterations, including point mutations and deletions, are among the most frequent genomic events in this tumor type, occurring in approximately 30%–60% of cases (1–3). Osteosarcoma is the most common primary malignant bone tumor in children and adolescents (4, 5). Despite marked molecular heterogeneity, standard treatment remains based on neoadjuvant chemotherapy, surgery, and adjuvant chemotherapy, resulting in an overall 5-year survival rate of approximately 60%–70% (6, 7). However, patients who respond poorly to standard therapy have substantially worse outcomes, with 5-year survival rates below 20% (5–8). In our previous multi-omics study, we identified RB1 as one of the most frequently altered genes in osteosarcoma and showed that molecular subtype information could inform stratified treatment strategies (9). RB1-deficient osteosarcoma is associated with aggressive disease behavior, reduced sensitivity to chemotherapy, and poor clinical outcome (10). Thus, defining actionable vulnerabilities created by RB1 loss may provide a rational framework for developing precision therapeutic strategies for this difficult-to-treat tumor state.

Direct pharmacological targeting of RB1 remains challenging because the RB1 protein lacks conventional small-molecule binding pockets, and therapeutic restoration of tumor-suppressor function is generally more difficult than inhibition of an oncogenic driver (11, 12). Synthetic lethality offers an alternative strategy by exploiting dependencies that arise specifically from cancer-associated genetic alterations. This approach can expand the spectrum of actionable targets, improve tumor selectivity, and potentially repurpose agents whose broad activity has limited their clinical development (13, 14). In RB1-deficient cancers, most synthetic-lethal strategies have focused on cell-cycle-associated vulnerabilities. For example, inhibition of Aurora kinase A or B has shown synthetic-lethal activity in RB1-deficient small-cell lung cancer models (15). However, therapies that further perturb cell-cycle machinery may have limited therapeutic windows and can produce substantial toxicity. Whether RB1 loss creates non–cell cycle dependencies that can be therapeutically exploited remains incompletely understood.

Proteostasis represents one such potential vulnerability. Valosin-containing protein (VCP/p97) is an evolutionarily conserved AAA+ ATPase that maintains protein homeostasis, organelle quality control, and genomic stability by facilitating the extraction and degradation of ubiquitinated substrates through multiple protein-quality-control pathways (16–18). Because tumor cells often experience elevated protein synthesis, metabolic stress, and organelle stress, they may become particularly dependent on VCP-mediated proteostasis. The endoplasmic reticulum (ER) is especially sensitive to perturbations in protein folding, and accumulation of misfolded or aberrant polypeptides activates adaptive stress programs, including the unfolded protein response, translational attenuation, and enhanced degradation of damaged proteins (19–23). Here, using osteosarcoma as a model of RB1-deficient malignancy, we performed high-throughput compound screening and identified VCP inhibition as a selective vulnerability associated with RB1 loss. We further show that this vulnerability is linked to glutamine-regulated VCP acetylation and activation of ER proteostasis stress. These findings uncover a metabolic–proteostasis dependency in RB1-deficient tumors and nominate VCP inhibition as a therapeutic strategy for this difficult-to-target tumor state.

## METHOD DETAILS

### Experimental model and study participant details

The study protocol was reviewed and approved by the Institutional Research Ethics Committee of Shanghai General Hospital, Shanghai Jiao Tong University School of Medicine (Approval No. 2021KY103). Prior to participation and sample collection, written informed consent was obtained from all participants or, where applicable, from their legal guardians. All research involving human subjects was carried out in compliance with internationally accepted ethical standards, including the principles outlined in the Declaration of Helsinki.

We established a cohort of RB1-deficient osteosarcoma patients, including 30 cases with RB1 deficiency and 30 cases without RB1 deficiency (a total of 60 osteosarcoma patients, **Supplementary Table 1**). All patients in this cohort received standardized treatment and surgery. The dataset includes information on age, sex, RB1 status, tumor location, pre-and post-neoadjuvant chemotherapy (preoperative) MRI imaging data, as well as metastasis and recurrence status.

We constructed a small-scale follow-up cohort of osteosarcoma patients, which included both recurrent and non-recurrent cases (a total of 27 patients as of 2023, re-OS mini-cohort 2023 edition, **Figure 3 and Supplementary Table 4**). All patients in this cohort underwent standardized treatment and surgery, including 12 non-recurrent and 15 recurrent cases. This includes age, gender, Enneking stage, tumor location, and VCP immunofluorescence (IF) intensity. Pathological diagnoses of all OS patients admitted to Shanghai General Hospital were independently examined by three pathologists. Clinical information and gene expression related to patient survival (GSE42352) for survival analysis were downloaded from Gene Expression Omnibus (GEO, https://www.ncbi.nlm.nih.gov/geo/) and Kaplan-Meier survival curves were compared using a two-sided log-rank test using R2 Genomics Analysis and Visualization Platform (R2, https://hgserver1.amc.nl/cgi-bin/r2/main.cgi).

**Figure 1.**
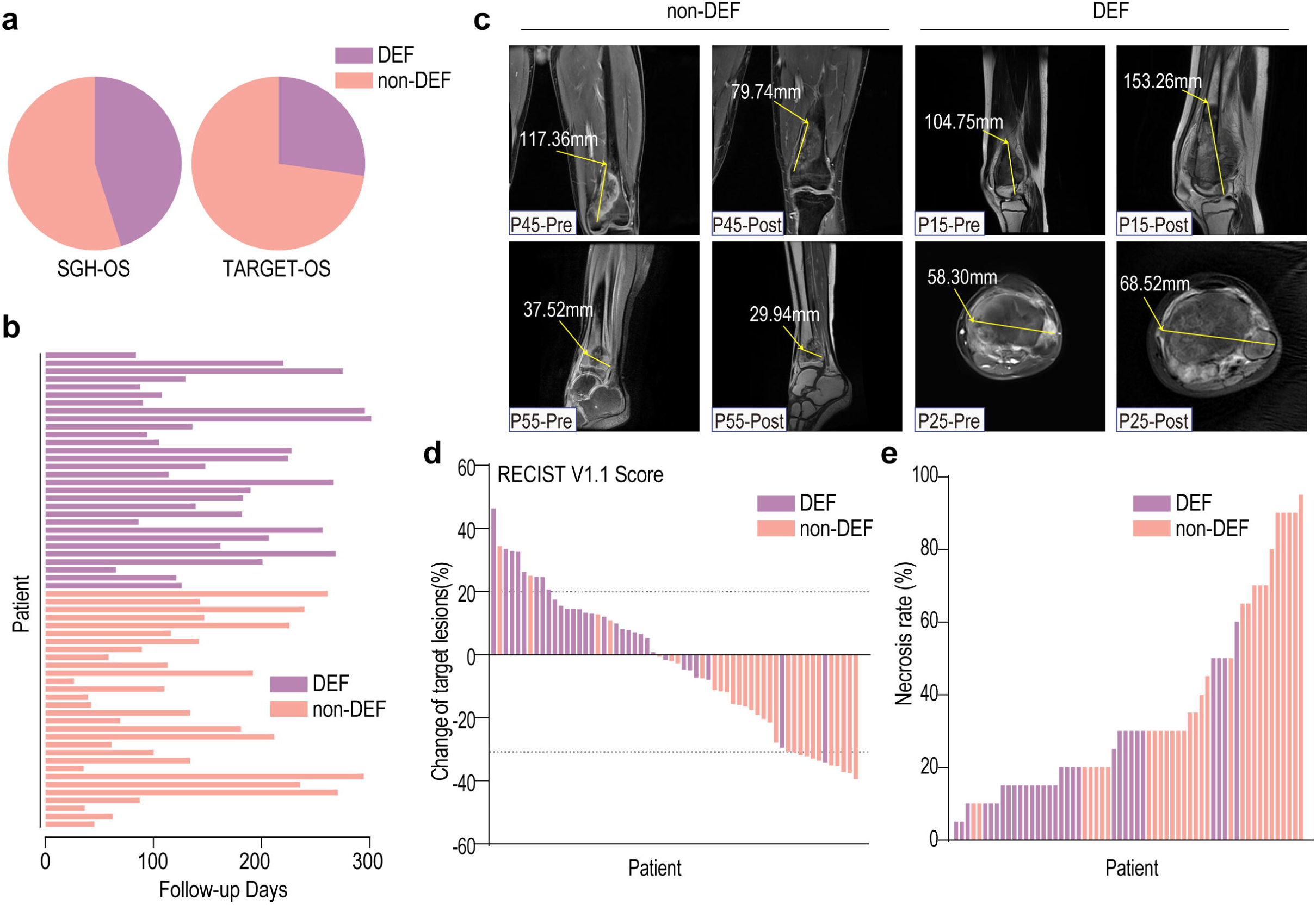
RB1-deficient osteosarcoma demonstrates high prevalence and poor neoadjuvant chemotherapy response. **(a)** Proportion of RB1-deficient patients in the SGH-OS and TARGET-OS cohorts, approximately 45% and 35%, respectively, as shown in pie charts. DEF indicates RB1-deficient, and non-DEF indicates RB1 non-deficient. **(b)** Follow-up duration for 60 clinically annotated osteosarcoma patients (30 DEF and 30 non-DEF). Each bar represents one patient. **(c)** Representative MRI scans from non-DEF (P45, P55) and DEF (P15, P25) patients before and after neoadjuvant chemotherapy, with maximum tumor diameter quantified on T1-weighted or T2-weighted sequences as appropriate. **(d)** Waterfall plot showing percentage change in target lesion size following neoadjuvant chemotherapy, assessed using RECIST v1.1 criteria. Each bar represents one patient. **(e)** Pathological necrosis rate (%) assessed in resected tumor specimens after neoadjuvant chemotherapy across the same patient cohort. Bars represent individual patients, and necrosis rate was independently evaluated by two musculoskeletal pathologists according to standard osteosarcoma grading criteria.

### Compound screens

RB1-deficient (shRB1) and control (shNC) HOS and MG63 cells were screened against a library of 2,097 small-molecule compounds (compound list provided in **Supplementary Table 2**). Cells were seeded at 800 cells per well in 50μL medium in 384-well plates, and 0.1μL of each 1 mM compound was transferred using a pin-tool system to achieve a final concentration of 2μM. Each plate contained 32 DMSO-only wells as negative controls. After 4 days of incubation, cell viability was measured using CellTiter-Glo (10μL/well; Promega #G7570). Raw luminescence values were normalized to the median of the DMSO control wells for each plate.

For each compound in each cell line, we calculated the difference in normalized viability between shRB1 and shNC cells, which served as a metric of RB1-selective drug sensitivity. These values—together with raw and normalized measurements—are provided in **Supplementary Table 3,** allowing full reproducibility of the compound-ranking procedure.

### Compound screens analysis

Cell viability data from the small-molecule compound screen were analyzed using custom R scripts and Excel. Raw luminescence values obtained from the CellTiter-Glo assay were normalized to the mean luminescence signal of DMSO-treated control wells within each experimental replicate.

Normalized viability values were calculated as:

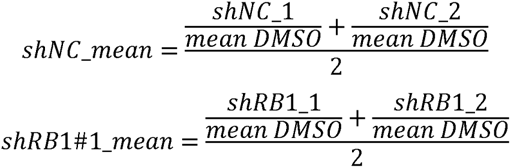

The raw screening data included HOS shNC_1, HOS shNC_2, HOS shRB1#1_1, and HOS shRB1#1_2 measurements. Normalized viability values included HOS shNC_mean, HOS shRB1#1_mean, log₂ (HOS shNC_mean), and log₂ (HOS shRB1#1_mean). RB1-selective sensitivity metrics included HOS shRB1#1_mean / HOS shNC_mean and log₂ (HOS shRB1#1_mean / HOS shNC_mean). Equivalent calculations were performed for MG63 cells. All raw screening data, normalized viability values, RB1-selective sensitivity metrics, and corresponding calculation formulas are provided in **Supplementary Table 3**.

To facilitate direct execution of the computational workflow, log₂ (HOS shNC_mean) and log₂ (HOS shRB1#1_mean) were extracted into **Supplementary Table 12**, whereas log₂ (MG63 shNC_mean) and log₂ (MG63 shRB1#1_mean) were extracted into **Supplementary Table 13** as standardized code-ready input files. **Supplementary Tables 12-13** were used as direct inputs for the R scripts generating the scatter and distribution plots shown in **Figure 2b-c**.

**Figure 2.**
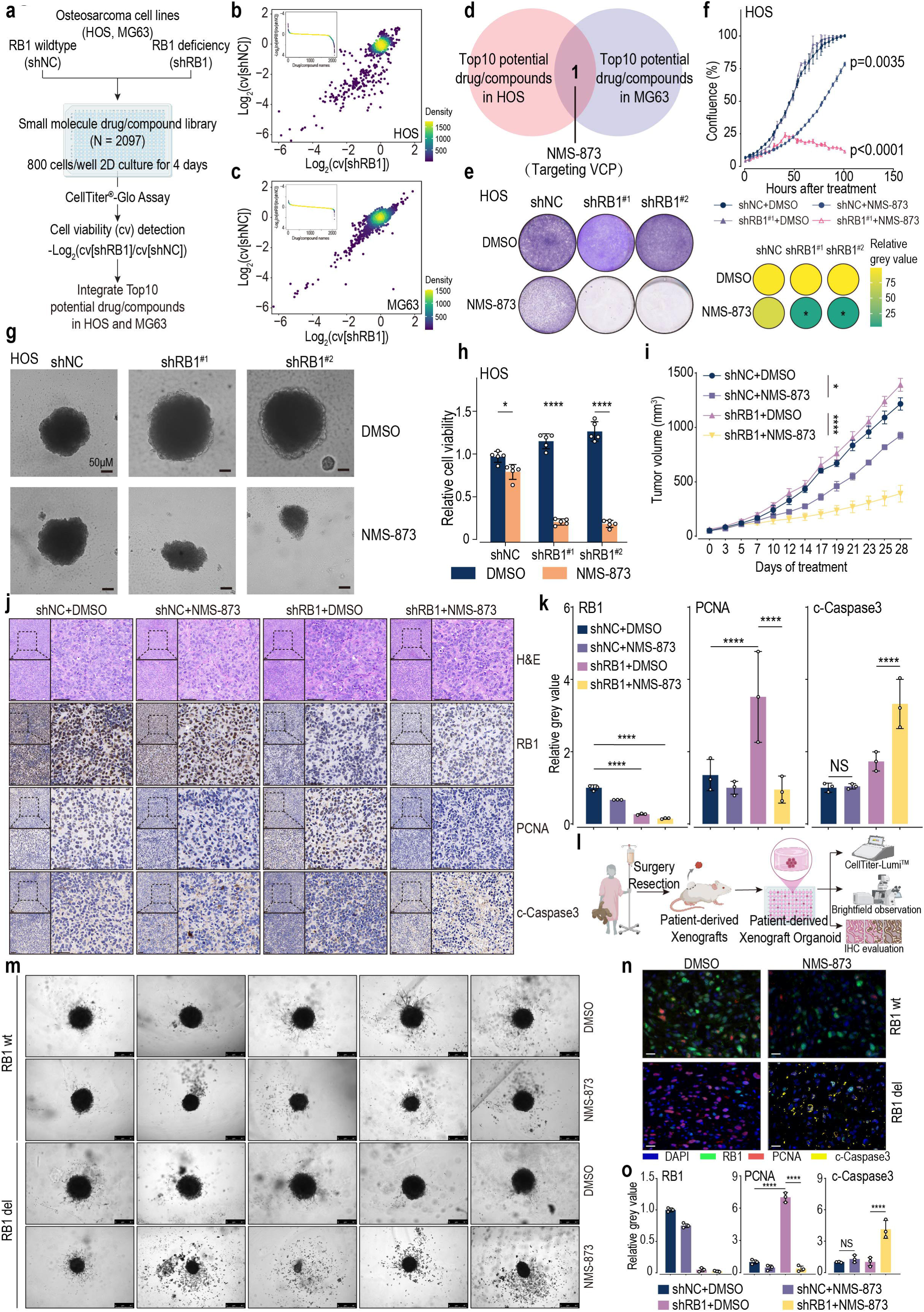
High-throughput compound screening identifies NMS-873 as a selective inhibitor of RB1-deficient osteosarcoma. Unless otherwise indicated, NMS-873 was used at 2μM for in vitro experiments and 20mg/kg for in vivo studies. **(a)** Workflow of high-throughput screening using a small-molecule compound library (N = 2,097) in RB1 wild-type (shNC) and RB1-deficient (shRB1) osteosarcoma cell lines (HOS and MG63). **(b, c)** Drug sensitivity distribution in HOS (b) and MG63 (c) cells based on the difference in normalized cell viability between shRB1 and shNC cells. **(d)** Venn diagram showing the overlap of the top 10 candidate compounds from both HOS and MG63 screens. NMS-873 was the only common hit. **(e)** Clonogenic assay validating the effect of NMS-873 in HOS cells with different RB1 statuses. Data represent three biological replicates. * indicates p < 0.05, two-way ANOVA with RB1 status and treatment as factors. **(f)** Incucyte live-cell imaging showing growth curves of HOS cells treated with NMS-873 or DMSO. Biological replicates: N = 3; analyzed by two-way ANOVA. **(g)** Bright-field images from 3D spheroid assays showing the effect of NMS-873 in HOS cells. Scale bar, 50μm. **(h)** Quantification of HOS cell viability in 3D spheroid assays. * indicates p < 0.05, **** indicates p < 0.0001; biological replicates: N = 5; two-way ANOVA. **(i)** Tumor growth in HOS RB1 wild-type (shNC) or RB1-deficient (shRB1) osteosarcoma mouse orthotopic tibial xenograft models after treatment with NMS-873 or DMSO. * indicates p < 0.05, **** indicates p < 0.0001; biological replicates: N = 5; two-way ANOVA. **(j)** Representative H&E and immunohistochemistry (IHC) staining for RB1, PCNA, and cleaved Caspase-3 in tibial xenograft models. Scale bar, 50μm. **(k)** Quantification of IHC staining. NS indicates not significant; **** indicates p < 0.0001; independent staining experiments (N = 3); two-way ANOVA. **(l)** Schematic of patient-derived xenograft and organoid model construction. **(m)** Bright-field images showing the effect of NMS-873 on tumor organoids derived from RB1 wild-type and RB1-deficient osteosarcoma. Scale bar, 500μm. Representative images are shown; quantitative viability measurements of the same organoid experiments are provided in **Supplementary** Figure 2d. **(n)** Immunofluorescence co-staining for RB1, PCNA, and cleaved Caspase-3 in tumor organoids. Scale bar, 20μm. **(o)** Quantification of immunofluorescence signal intensity. NS indicates not significant; *** indicates p < 0.001, **** indicates p < 0.0001; independent staining experiments (N = 3); two-way ANOVA.

**Figure 3.**
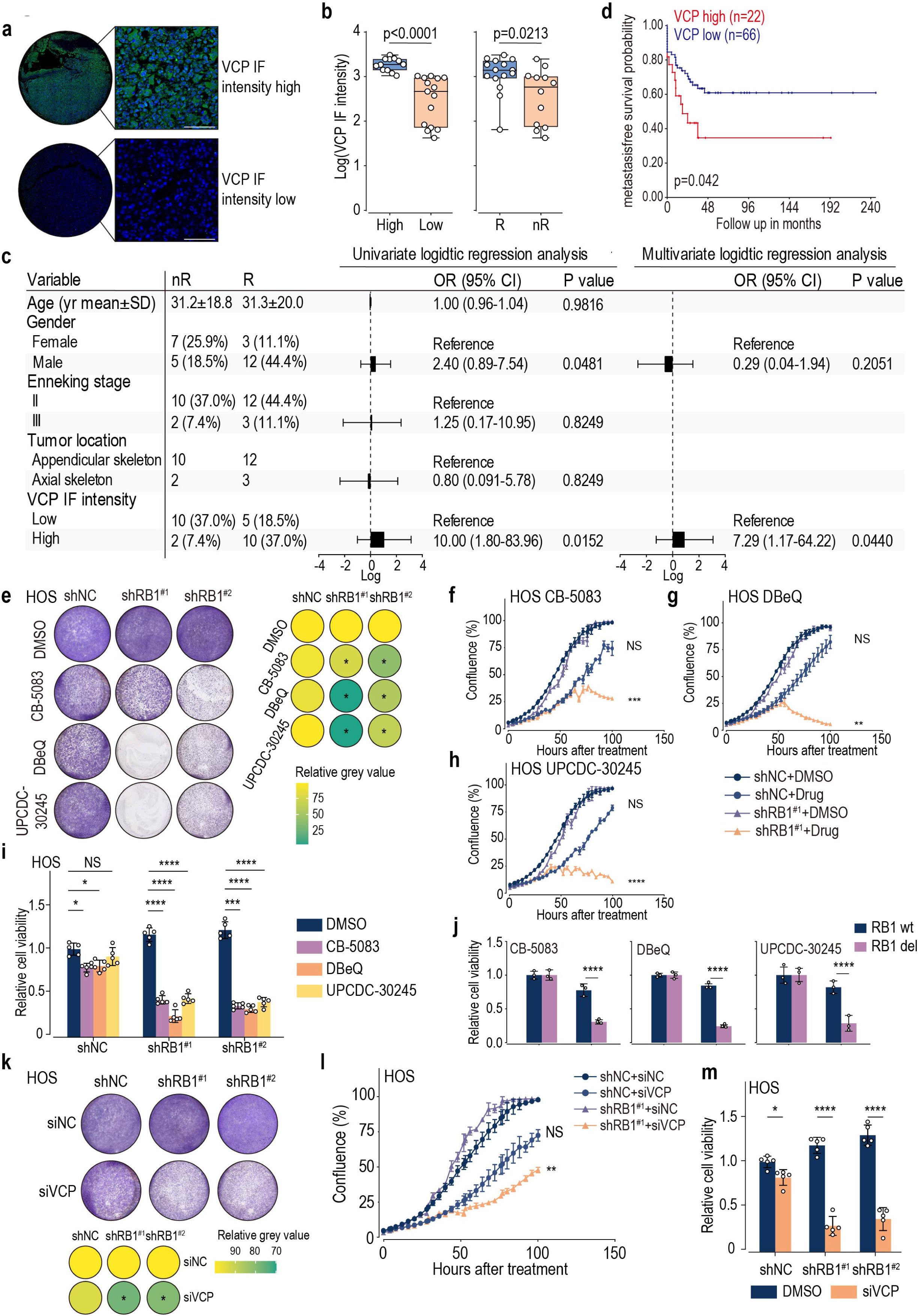
Targeting VCP selectively inhibits RB1-deficient osteosarcoma. **(a)** Immunofluorescence staining of VCP expression in osteosarcoma tissue microarrays. Scale bar, 100μm **(b)** Quantification of VCP fluorescence intensity. Samples were classified into high (N = 12) and low (N = 15) VCP expression groups. Comparison of VCP levels between patients with disease recurrence (R, N = 15) and non-recurrence (nR, N = 12). Statistical analysis was performed using a t-test. **(c)** Summary of clinical characteristics of 27 patients from the tissue microarray cohort, including age, sex, Enneking stage, tumor location, VCP expression group, and disease progression status. Univariate and multivariate logistic regression analyses were performed to evaluate factors associated with disease progression. **(d)** Kaplan–Meier survival analysis of metastasis-free survival in osteosarcoma patients with high (N = 22) or low (N = 66) VCP expression (GSE42352 dataset). p = 0.042. **(e)** Clonogenic assays showing the effects of VCP inhibitors CB-5083, DBeQ, and UPCDC-30245 in HOS cells. Quantification based on three biological replicates. * indicates p < 0.05; analyzed by two-way ANOVA. Cells were treated with CB-5083 (0.5μM), DBeQ (2μM), or UPCDC-30245 (2μM) for 5 days. **(f–h)** Incucyte live-cell imaging showing the effects of CB-5083 (f), DBeQ (g), and UPCDC-30245 (h) on HOS cell proliferation. NS indicates not significant; ** indicates p < 0.01, *** indicates p < 0.001, **** indicates p < 0.0001. Biological replicates: N = 3; analyzed by two-way ANOVA. Cells were treated with CB-5083 (0.5μM), DBeQ (2μM), or UPCDC-30245 (2μM) for 100 hours. **(i)** Cell viability assessment from 3D spheroid assays in HOS cells treated with CB-5083, DBeQ, or UPCDC-30245. NS indicates not significant, * indicates p < 0.05, *** indicates p < 0.001, **** indicates p < 0.0001. Biological replicates: N = 5; two-way ANOVA. Cells were treated with CB-5083 (0.5μM), DBeQ (2μM), or UPCDC-30245 (2μM) for 100 hours. **(j)** Cell viability of osteosarcoma organoids treated with CB-5083, DBeQ, or UPCDC-30245. **** indicates p < 0.0001; technical replicates: N = 3; two-way ANOVA. Cells were treated with CB-5083 (0.5μM), DBeQ (2μM), or UPCDC-30245 (2μM) for 100 hours. **(k)** Clonogenic assays showing the effect of VCP knockdown (siVCP) in HOS cells. Quantification based on three biological replicates. * indicates p < 0.05; two-way ANOVA. **(l)** Incucyte proliferation curves of HOS cells with siVCP knockdown. NS indicates not significant, ** indicates p < 0.01. Biological replicates: N = 3; two-way ANOVA. **(m)** Cell viability from 3D spheroid assays after siVCP treatment in HOS cells. * indicates p < 0.05, *** indicates p < 0.001, **** indicates p < 0.0001. Biological replicates: N = 5; two-way ANOVA.

Compounds were subsequently ranked in ascending order according to the RB1-selective sensitivity metrics: log₂ (HOS shRB1#1_mean / HOS shNC_mean) and log₂ (MG63 shRB1#1_mean / MG63 shNC_mean). The top 10 compounds from each cell line were identified independently, and their intersection was visualized using a Venn diagram to generate **Figure 2d**.

The complete analysis workflow and annotated code are available through the verified Code Ocean compute capsule: https://codeocean.com/capsule/4710920/tree/v1

### Cell lines, antibodies, and reagents

The human OS cell lines HOS (Cat#CRL-1543; CVCL_0312), MG-63 (Cat#CRL-1427; CVCL_0426), U2-OS (Cat#HTB-96; CVCL_0042) and the human kidney cell line HEK-293T were cultured in high glucose Dulbecco’s Modified Eagle Medium (DMEM; Gibco, USA) supplemented with 10% fetal bovine serum (FBS; Wisent, Canada). Cell line authentication was performed on cells that used for in vitro and in vivo studies using Short Tandem Repeat (STR) DNA profiling and all cell lines were obtained from the Chinese Academy of Medical Sciences were preserved from the Shanghai Bone Tumor Institute (Shanghai, China), Shanghai General Hospital. Reagents used in the study included NMS-873 (#S7285, Selleck, USA, RRID:), CB-5083 (#S8101, Selleck, USA), DBeQ (#S7199, Selleck, USA), UPCDC-30245 (#HY-123636, MCE, USA), Thapsigargin (#S7895, Selleck, USA), Tunicamycin (#S7894, Selleck, USA), RH01687 (#S3461, Selleck, USA), RH01386 (#HY-124771, MCE, USA). The antibodies used in this study were purchased as follows: anti-GAPDH (#ab181602, 1:1000, RRID: AB_2630358), anti-IRE1 (#ab37073, 1:2000, RRID: AB_775780), and anti-ATF6 (#ab37149, 1:1000, RRID: AB_725571) were all purchased from Abcam. Anti-XBP1 (#40435, 1:1000, RRID:AB_2891025), anti-PERK (#5683, 1:1000), anti-EIF2α (#5324, 1:1000), anti-HSPA5(#3177, 1:1000), anti-VCP (#2648, 1:1000, Western-blotting used), anti-Acetylated-Lysine(#9441, 1:1000), anti-RB1(#9309, 1:2000, Western-blotting used; 1:1000, IHC-staining used) and anti-CHOP (#2895, 1:1000) were all purchased from Cell Signaling Technology. anti-VCP (#GB121155-100, 1:100, IF-staining used), anti-PCNA (#GB11010-100,1:200, IF-staining used) and anti-Cleaved-Caspase-3 (#GB11532-100, 1:500, IF-staining used) were all purchased from Servicebio.

All cell lines underwent STR profiling and were tested for Mycoplasma monthly using the Mycoplasma Detection Kit (catalog no. rep-mys-20, InvivoGen) and used within 10 passages.

### Mouse xenografts of osteosarcoma

6-8-week-old female Nude mice (Strain: BALB/cJGpt-foxn1nu/Gpt; Charles River, USA/China) (RRID: IMSR_GPT: D000521) were used in this study. All mice were maintained under SPF conditions in a controlled environment of 20-22℃, with a 12/12h light/dark cycle, 50-70% humidity. All animal experiments were performed by following the protocols in accordance with the guidelines of Laboratory Animal Center of Shanghai General Hospital. The Clinical Center Laboratory Animal Welfare & Ethics Committee of Shanghai General Hospital, Shanghai Jiao Tong University School of Medicine, approved all animal protocols used in this study (IACUC: 2023AW67). The OS mouse xenograft was developed by intramedullary injection with OS cells (5×10^5^ cells in 25μL PBS supplemented with 10% FBS) into the marrow space of the proximal tibial or the lateral abdominal region of nude mice with a 27-gauge needle, as our previous studies(24). OS progression in the mice was monitored by measuring the tumor volume, calculated by the following equation: Volume = Length×Width²/2. The tumor burden was monitored by following the tumor volume and the maximal tumor size is 2 cm, which was permitted by the Clinical Center Laboratory Animal Welfare & Ethics Committee of Shanghai General Hospital. The maximal tumor size was confirmed to be not exceeded in our study and the survival curves were produced according to the survival of mice and the time to reach maximum tumor size.

### Construction and cultivation of organoid models

Tumor of patient-derived xenograft (PDX) tissues were digested into single cells and allowed to self-assemble to construct organoid models. Culture methods from our previous research were applied for cultivation and drug treatment, and cell viability assays were performed after approximately 7 days. The specific construction and cultivation methods are described in our earlier studies(25).

### Immunohistochemistry (IHC), immunofluorescence (IF) and hematoxylin and eosin (H&E) stainings

IHC staining was performed on representative tissue sections from formalin-fixed and paraffin-embedded tissue blocks from human OS TMA and the mouse tibia orthotopic tumor models using the mentioned antibodies at the indicated concentrations. IF staining was performed on representative tissue sections from formalin-fixed and paraffin-embedded tissue blocks from human OS TMA and the mouse tibia orthotopic tumor models using the mentioned antibodies at the indicated concentrations. The hematoxylin and eosin (H&E) staining of human osteosarcoma tissue microarrays (OS TMA) and the mouse tibia orthotopic tumor models was conducted following the methods used in our previous study 27. For live-cell aggregate IF staining, refer to the PROTEOSTAT® Aggresome Detection Kit (#ENZ-51035-K100) manual for detailed instructions.

### Western blotting

Proteins were extracted by using radio immunoprecipitation assay (RIPA) lysis buffer (Beyotime, China) for total protein and Histone Extraction Kit (#ab113476, Abcam, USA) for histones according to the manufacturer’s protocol. The extracted protein was quantified using a Pierce BCA Protein Quantification Kit (#23325, Thermo Fisher Scientific, USA) and measured with a SpectraMax M3 Microplate Reader (Molecular Devices, USA). Proteins were separated by sodium dodecyl sulfate polyacrylamide gel electrophoresis (SDS-PAGE) and transferred to 0.45um polyvinylidene fluoride (Millipore, USA) using Mini-PROTEAN Tetra Vertical Electrophoresis Cell electrophoresis chamber and Mini Trans-Blot Module for tank transfer system with PowerPac HV Power Supply (Bio-Rad, USA, RRID:SCR_008426). The membrane was blocked using 5% nonfat dry milk in Tris-buffered saline solution containing 0.1% Tween-20 (TBS-T) for 1h at room temperature and then probed with specific primary antibodies overnight at 4℃. Subsequently, the membranes were washed three times with TBS-T for 10min each, followed by incubating with HRP-conjugated secondary antibodies for 1h at room temperature. Actin and GAPDH were used as the protein loading control. Protein signals were developed with SuperSignal West Femto Maximum Sensitivity Substrate (Thermo Fisher Scientific, USA) and imaged using chemiluminescence imaging system Amersham Imager 600 (GE Healthcare, USA, RRID:SCR_021853) and Tanon 5200 (Tanon, China, RRID:SCR_027114).

### Immunoprecipitation and immunoblotting

Proteins were extracted from cultured cells using a modified buffer (50mM Tris-HCl (pH 7.4), 150mM NaCl, 1mM EDTA, 10% glycerol and 1% NP-40) supplemented with protease and phosphatase inhibitor cocktail (Thermo, #78441) and PMSF (Thermo, #36978), followed by immunoprecipitation and immunoblotting with the corresponding antibodies. Cell lysates were collected after centrifugation to remove cell debris. For exogenous immunoprecipitation, cell lysates were incubated with FLAG beads (Sigma, #M8823) at 4°C overnight, and the beads were boiled after extensive washing. For endogenous immunoprecipitation, cell lysates were incubated with the indicated antibodies at 4°C overnight, and the immunoprecipitate was incubated with protein A/G agarose beads (Thermo, #20422) for 2-3h followed by washing. Protein samples were separated by SDS-PAGE, transferred onto PVDF membrane (Millipore), and probed with the indicated antibodies.

### Real-Time PCR

For RT-PCR, EZ-press RNA purification kit (EZBioscience, Roseville, United States) was used to collect the purified total RNA in cultured cells. Then, a color-reverse transcription kit (EZBioscience, Roseville, United States) was applied to remove the genomic DNA (gDNA) and the reverse transcription of RNA to get the cDNA being performed. 2 × color SYBR green qPCR master mix (EZBioscience, Roseville, United States) was used for RT-PCR on QuantStudio 6 Flex RT-PCR systems (Thermo Fisher Scientific Inc., United States). Ct values were obtained from technical triplicates and averaged for each biological replicate. Relative mRNA expression values shown in the figures were calculated using the 2–ΔΔCt method, with GAPDH as the internal control and the shNC+DMSO group as the calibrator. Error bars represent the variability across biological replicates.

### Primer Design

Primers for the target and reference genes were designed using the Primer3 software and synthesized by Sigma Aldrich (USA). The sequences of the primers (Forward primers) used provided in **Supplementary Table 14.**

### Colony formation

To assay cell growth, OS cells were washed twice with PBS and plated onto 6-well cell culture plates at a density of 30,000 cells per well. OS cells were treated with or without drugs for 1 week, and the cell colonies were stained using crystal violet staining solution (Beyotime, China) according to the manufacturer’s instructions.

### Incucyte cell proliferation assay, cell viability assay and Caspase-3/7 apoptosis assay

Indicated cell lines were seeded into 96-well plates at a density of 1,000–2,000 cells per well, depending on the growth rate and the design of the experiment. About 24 hours later, the indicated compounds were added at the indicated concentrations. Cells were imaged every 4 hours using the Incucyte ZOOM (Essen Bioscience). Phase-contrast images were analyzed to detect cell proliferation on the basis of cell confluence. For the cell viability assay, CellTiter-blue (Promega, #G8081) was added to the medium following the manufacturer’s instructions. For the cell apoptosis, Caspase-3/7 apoptosis-assay reagent was added to the culture medium, and cell apoptosis was analyzed on the basis of fluorescent staining of apoptotic cells.

### Glutamine-restricted and glucose-restricted culture

Custom media without D-glucose, sodium pyruvate, and L-glutamine (#X070M1, BasalMedia, China) was used. For glutamine restriction, 4.5g/L glucose and 1 mM sodium pyruvate were added to the custom media. For glucose restriction, 4mM glutamine was added to the custom media.

### LC-MS for targeted metabolomics

For extracting metabolites from cells, HOS cells (0.5-1×10^7^ per sample) were washed three times with ice-cold normal saline (0.9% NaCl) and metabolism was quenched by adding 4 mL of 80% methanol (cooled to -80°C). After 20 min of incubation on dry ice, cells were scraped and centrifuged at 14,000g for 10 minutes at 4°C. Insoluble pellets were re-extracted with 500μL of prechilled 80% methanol on dry ice. The supernatants from two rounds of extraction were combined. To precipitate proteins, 3 mL of chloroform was added into the tubes, and vortexed for 1 min, then centrifuged at 14,000g for 5min at 4°C. Supernatants were thoroughly lyophilized (FreeZone 6 Liter, Labconco, USA) and reconstituted in 50μL of methanol-water (1:1, v/v) just prior to measurement. Metabolites were analyzed using a Vanquish UPLC system (Thermo Scientific) & Q Exactive plus Mass spectrometer (Thermo Scientific) coupled with hydrophilic interaction chromatography (HILIC). LC separation was on a SeQuant ZIC-HILIC column (100 mm × 2.1mm i.d., 3.5µm) (Merck) using a gradient of solvent A (50mM ammonium formate in water) and solvent B (acetonitrile). The gradient was 0-10min, 90-50% B; 10-12min, 50-90% B; 12-15min, 90% B; and the column was maintained at 45°C. The flow rate was 0.4 mL/min. The MS scans were in both negative and positive ion modes with a resolution of 140,000 at m/z 200. The scan range was m/z 60– 900. Lock masses were used and the mass accuracy obtained for all metabolites was below 5 p.p.m. Data were analyzed using TraceFinder 5.0 (Thermo Scientific). For targeted metabolomics, the extraction solvent contained 13C6 glucose-6-phosphate (10μM), d4-succinate (10μM), and 2-Cl-phenylalanine (10μM) as internal standards. The data were normalized to protein concentration.

### Determination of extracellular fluxes

The glucose level in the culture medium was measured by HPLC (model 1260, Agilent, Santa Clara, USA) using a cation-exchange column (HPX-87H, Bio-Rad, Hercules, CA) and a differential refractive index (RI) detector. A mobile phase of 5mM H2SO4 at 0.5mL/min flow rate was used and the column was operated at 60°C. Glutamine level in the culture medium was measured by ACQUITY Ultra Performance LC systems using the Waters AccQ. Tag Amino Acid Analysis Method (Waters, USA). The specific growth rate, and specific uptake rates of glucose and glutamine were determined by solving the equations as described28. The data were normalized to cell number.

### Stable gene overexpression and shRNA Knock-Down construction

VCP overexpression using plasmid (1:5’-CGCGAATTCGAAGTATACCTCGAGGCCACCATGGCTTCTG-3’), and constitutive RB1 knock-down using shRNA hairpins targeting RB1 (#1: 5’-CCACATTATTTCTAGTCCAAA-3’, #2: 5’-CGCGTGTAAATTCTACTGCAA-3’).

### Stable gene overexpression and shRNA Knock-Down construction

For RB1 knock-down (shRB1) or VCP over-expression (OE VCP), Lentivirus production was obtained from PEI transfection reagents (#26406, Polyseciences, USA) of HEK-293T cells with co-transfection of the packaging vectors pspAX2 (RRID: Addgene_12260) and pMD2.G (RRID: Addgene_12259) along with the gene delivery vector. Viral supernatants were collected 72h after transfection, underwent ultracentrifugation at 20,000rpm for 25h at 4℃ to concentrate, and the virus pellets were resuspended in PBS. For infection, the viral pellets were added to cells in a dropwise manner in the presence of polybrene (10μg/ml). After 48h, medium containing the lentivirus was replaced and infected cells were selected by addition of puromycin (2ug/ml).

### Small interfering RNA (siRNA) interference experiment

On the day before transfection, 0.5 – 2×10^5^ cells were seeded in 400μL of antibiotic-free medium to achieve a cell density of 30% -50%. The siRNA (Genomeditech, Shanghai, China) was diluted (final concentration at 50nM) in 50μL of Opti-MEM (Gibco, USA) and gently mixed by pipetting. Separately, 1.0μL of Lipofectamine™ 3000 (Thermo Fisher Scientific, USA) was diluted in 50μL of Opti-MEM, gently mixed by pipetting, and allowed to stand at room temperature for 5 minutes. The transfection reagent and siRNA dilution were then combined to form the transfection complex, gently mixed by pipetting, and incubated at room temperature for 20 minutes. The transfection complex was gently added dropwise into the cell well plate and mixed by cross-pipetting. The cells were incubated at 37°C with 5%CO_2_ for 18h to 48h, with the medium replaced with complete medium 4h to 6h after transfection. The silencing efficiency of the siRNA was validated using Western-blot (WB).

### Protein extraction and trypsin digestion

20µg protein for each sample were mixed with 2X loading buffer respectively and boiled for 5min. The proteins were separated on SDS-PAGE gel (constant current 120 V, 60 min). Protein bands were visualized by Coomassie Blue R-250 staining. Target Protein bands were collectted and digestion was performed by trypsin. According to the experimental design, samples were obtained from target protein to collect gel points. After the gel points underwent destaining and fragmentation, a solution containing 20ng/uL trypsin in 50mM NH_4_HCO_3_ was added. Enzymatic digestion took place in a 37°C incubator for 16 hours, followed by the extraction and freeze-drying of peptide extracts. Prior to analysis, the samples were resolubilized in a 0.1% formic acid solution and underwent peptide quantification before being subjected to mass spectrometry analysis.

### Histone Extraction

Histones were extracted from cultured osteosarcoma cells using the Abcam Histone Extraction Kit (#ab113476) with the following streamlined procedure: Adherent cells (∼1×10^7^) were harvested by trypsinization, washed once in cold PBS, and pelleted (1,000g, 5min, 4°C). Pellets were resuspended in Pre-Lysis Buffer (kit) at 1×10^7^cells/mL, incubated on ice for 10min with occasional mixing, then spun (3,000g, 5min, 4°C). The supernatant was discarded. Pellets were lysed in Acid Extraction Buffer (200µL per 1×10^7^cells) on ice for 30min, vortexing briefly every 10min. Samples were centrifuged at 12,000g for 5min (4°C), and the acid-soluble supernatant containing histones was collected. Supernatants were neutralized by adding Balance-DTT Buffer (0.3×volume), then protein concentration was measured against BSA standards. Histone extracts were aliquoted and stored at –80°C until use.

### 3D cell viability assay

Cell viability was measured using a CellTiter-Lumi™ luminescent cell viability assay kit (#C0065S, Beyotime). The preparation and protocol of the cell viability assay kit were performed according to the Beyotime official protocol and were abbreviated as follows. The cell culture plates were allowed to equilibrate at room temperature for 10min (not exceeding 30min). Next, 100μL of CellTiter-Lumi™ luminescent assay reagent per well was added to a 96-well plate. The cells were gently shaken at room temperature for 2min to facilitate cell lysis. The cells were incubated at room temperature for 10min to stabilize the luminescence signal. Luminescent detection was performed using a SpectraMax M3 Microplate Reader (Molecular Devices). Relative cell viability was calculated directly from the luminescent readings or ATP content was determined using an ATP standard curve to assess relative cell viability based on ATP levels. Nuclease-free ATP was purchased from Beyotime (#D7378, Beyotime).

### Nano-LC-MS/MS analysis

For the proteome profiling samples, peptides were analyzed on an OE480 Hybrid Quadrupole-Orbitrap Mass Spectrometer (Thermo Fisher Scientific) coupled with a high-performance liquid chromatography system (EASY nLC 1200, Thermo Fisher Scientific). Dried peptide samples were re-dissolved in Solvent A (0.1% formic acid in water) were loaded onto a 2-cm self-packed trap column (100μm inner diameter, Dr. Maisch GmbH) using Solvent A and separated on a 150-μm-inner-diameter column with a length of 15cm (1.9μm ReproSil-Pur C18-AQ beads, Dr. Maisch GmbH) over a 75-min gradient (Solvent A: 0.1% formic acid in water; Solvent B: 0.1% formic acid in 80% ACN) at a constant flow rate of 600 nL/min (0– 75min, 0min, 4% B; 0-10min, 4–15% B; 10–60min, 15–30% B; 60–69min, 30–50% B; 69–70min, 50–100% B; 70–75min, 100% B). Eluted peptides were ionized at 2.4kV and introduced into the mass spectrometer. Mass spectrometry was performed in data-dependent acquisition mode. For the MS1 Spectra full scan, ions with m/z ranging from 300 to 1,400 were acquired by an Orbitrap mass analyzer at a high resolution of 120,000. The automatic gain control (AGC) target value was set to 3E+06. The maximal ion injection time was 80 ms. MS2 spectral acquisition was performed in a rapid speed mode with 1s cycle time. Precursor ions were selected and fragmented with higher energy collision dissociation (HCD) with a normalized collision energy of 30%. Fragment ions were analyzed by an Orbitrap mass analyzer at a resolution of 7,500, with an AGC target at 5E+04. The maximal ion injection time of MS2 was 22ms. Peptides that triggered MS/MS scans were dynamically excluded from further MS/MS scans for 12 s. The coefficient of variation (CV) values for FAIMS were -45V and -65V. Nano-LC–MS/MS analysis was performed by iProteome Biotechnology Co., Ltd, Shanghai.

### Nano-LC–MS/MS data processing

The original data of mass spectrometry analysis were RAW files, and iProteome one-stop data analysis cloud platform was used for qualitative and quantitative analysis. Data processing and analysis was performed by iProteome Biotechnology Co., Ltd, Shanghai.

### Preparation and analysis of bulk RNA sequencing (RNA-seq)

Total RNA was isolated using TRIzol reagent (Invitrogen) following the manufacturer’s protocol. RNA quality and purity were evaluated through agarose gel electrophoresis and measured using a Nanodrop spectrophotometer (Thermo Fisher Scientific). Additionally, RNA purity was verified with a Kaiao K5500 Spectrophotometer (Kaiao, China), and its integrity and concentration were determined using the RNA Nano 6000 Assay Kit on the Bioanalyzer 2100 system (Agilent Technologies, USA). For each sample, 2 µg of total RNA was used to prepare sequencing libraries. Library construction was carried out using the NEBNext Ultra RNA Library Prep Kit for Illumina (NEB, USA), with unique index codes incorporated to distinguish individual samples. The indexed libraries were clustered on a cBot system using the HiSeq PE Cluster Kit v4-cBot-HS (Illumina, USA), following the supplier’s guidelines. Sequencing was performed on the DNBSEQ-T7 platform (RRID:SCR_017981, BGI, China) at Wuhan Benagen Technology Co., Ltd. (Wuhan, China), generating 150 bp paired-end reads.

The raw sequencing data were initially assessed using FastQC to remove adapter sequences and low-quality reads. Cleaned paired-end reads were then mapped to either the human reference genome (GRCh38) or mouse reference genome (mm10) using STAR (RRID:SCR_004463, v2.7.6a) (26). Only reads with high mapping quality (MAPQ > 30) and alignment to genomic exons were retained, and gene-level read counts were obtained using featureCounts(27) (referencing Ensembl 93 for GRCh38 or mm10). Differential expression analysis was conducted with the R package DESeq2 (RRID:SCR_015687)(28), employing the lfcShrink function to estimate fold changes. Genes were classified as significantly differentially expressed if they met the criteria: fold change ≥ 2.00, posterior probability ≥ 0.80, and false discovery rate (FDR) < 0.05. Functional enrichment analysis of differentially expressed genes was carried out using Gene Ontology (GO; http://geneontology.org/) through the clusterProfiler R package (v3.8)(29). Gene set enrichment analysis (GSEA; RRID:SCR_003199, http://www.gsea-msigdb.org/gsea/)(30) was performed using the GSEAPreRanked function, ranking genes by DESeq2-derived fold change, applying 1000 permutations, and considering gene sets significant when the normalized P value was < 0.05. Gene set variation analysis (GSVA) scores were computed for each sample using the GSVA R package, reflecting the relative activity of pathways across samples. These GSVA scores were subsequently analyzed to identify differentially modulated pathways under varying conditions. Gene sets were sourced from the Molecular Signatures Database (MSigDB, RRID:SCR_016863, https://www.gsea-msigdb.org/gsea/msigdb/)(31, 32). Prior to GSVA, the expression data underwent preprocessing and normalization. Finally, the calcPhenotype function (RRID:SCR_023872) was applied to integrate the RNA-seq profiles from the TARGET-OS cohort with the GDSC2 database (https://www.cancerrxgene.org/) to generate a drug response prediction matrix.

### Analysis of single-cell RNA sequencing (scRNA-seq)

The single-cell RNA sequencing (scRNA-seq) data used in this study comprised previously published data (n = 16), including 6 individuals from Liu et al. (GSE162454)(33) and 10 individuals from Zhou Y et al. (GSE152048)(34). The raw data were processed using the Seurat package (v4.0, http://satijalab.org/seurat/) and analyzed in R software (v4.2.0). The Seurat object contained gene expression data for each sample and was imported using the Read 10× function. We filtered out cells with fewer than 200 or more than 10000 detected genes, or with mitochondrial gene content exceeding 10% of total expressed genes. Data normalization was performed using log normalization with the default scaling factor. Subsequently, the highly variable genes (HVGs) were identified from the normalized expression matrix, centered, and scaled before conducting principal component analysis (PCA). Single-cell data integration and analysis were performed using the canonical correlation analysis (CCA) method in Seurat. Clustering analysis was conducted based on the integrated joint embedding data, and the Uniform Manifold Approximation and Projection (UMAP) method was used for visualization. Cell cluster annotation was performed using the Wilcoxon rank-sum test in Seurat, with Bonferroni correction applied to identify differentially expressed genes (DEGs) with high discriminatory ability. Cell subpopulations were annotated based on these DEGs. The analysis of cell receptor-ligand interactions was conducted using the iTALK package in R (https://github.com/Coolgenome/iTALK)(35). Given that osteosarcoma arises from the mesenchymal lineage, malignant tumor cells often resemble mesenchymal stromal cells (MSCs) in their transcriptomic profiles. Therefore, conventional tumor cell markers such as EpCAM are not applicable. In this study, we adopted a classification approach based on transcriptomic similarity and canonical lineage markers, which yielded mesenchymal, myeloid, lymphoid, and endothelial populations. This clustering strategy is consistent with previous osteosarcoma single-cell studies(36, 37). To distinguish malignant from non-malignant MSC subpopulations, we further examined proliferation signatures (*MKI67*, *PCNA*) and copy number variation (CNV) scores inferred by the inferCNV method.

### Quality control steps in scRNA-seq analysis

Mitochondrial gene content (percent.mt): For each sample, we used the PercentageFeatureSet function to calculate the mitochondrial gene content using the mitochondrial gene pattern “MT-”. Cells with mitochondrial gene content exceeding 10% were excluded, as this typically indicates that the cells are under stress or undergoing apoptosis. Gene count and mitochondrial proportion filtering: Cells were filtered based on the number of genes detected (200–10,000 genes) and mitochondrial gene content (below 10%). This step helps ensure that only viable, non-apoptotic cells are retained for analysis. Red blood cell gene expression (percent.HB): We calculated the expression of red blood cell genes (e.g., HBA1, HBB) and excluded cells with more than 1% red blood cell gene expression. This step removes cells that may be contaminated by red blood cells. Doublet detection and removal: Doublets were identified using the DoubletFinder package. The expected doublet proportion was set to 7.5% (0.075 * number of cells), and principal component analysis (PCA) with 30 principal components was used to detect doublets. Cells predicted to be doublets were removed from the dataset. Data normalization and variable gene selection: Data were normalized using the NormalizeData function, followed by detection of variable genes using the FindVariableFeatures function. We then performed data scaling using ScaleData and conducted PCA analysis on the normalized data.

### Bioinformatics and statistical analysis

For pathway analyses, over-representation enrichment was performed using all detected genes in each dataset as the background universe, with background gene sets provided in **Supplementary Table 6** (single-cell data) and **Supplementary Table 7** (MG63 RNA-seq). In contrast, pathway associations derived from the SGH-OS and TARGET-OS cohorts were based on correlation analysis with VCP expression and therefore did not require a background gene universe (**Supplementary Table 8-9**).

## QUANTIFICATION AND STATISTICAL ANALYSIS

Transcriptomes of patients in The Cancer Genome Atlas (TCGA, RRID: SCR_003193) were downloaded from the Genomic Data Commons (https://portal.gdc.cancer.gov/). The transcriptome of tumor cell lines was downloaded from the Cancer Cell Line Encyclopedia (CCLE) (https://sites.broadinstitute.org/ccle, RRID: SCR_013836) database. All the data were processed by R version 4.3.2 and some visualizations were performed using Posit (also RStudio, The Open-Source Data Science Company, RRID: SCR_000432). The interaction between VCP protein and NMS-873 was based on Pan M et al.(38) and visualized by PyMOL (version 2.4, RRID: SCR_000305). VCP acetylated proteome was according to Mori-Konya C et al.(39). The acetylation sites of VCP were predicted using the Group-based Prediction System version 6.0 (GPS, https://gps.biocuckoo.cn/, RRID:SCR_016374). Details of statistical analyses of the various experiments are described in the relevant methods section. If not specified, statistical analysis was carried out using GraphPad Prism 9 software (GraphPad Software, USA, RRID: SCR_000306). After confirming that values followed a normal distribution, two-tailed Student’s t test was applied to determine the significance of differences between two groups of independent samples. Spearman’s correlation analysis was performed to determine the correlation between two group of variables. The image was subjected to gray value analysis using ImageJ software (Thermo Fisher Scientific, USA, RRID:SCR_003070). The survival rate was analyzed using the mentioned Kaplan-Meier method. A p value < 0.05 was considered statistically significant. Details of the data points shown were described in the respective figure legends. All schematic diagrams were created using BioRender (BioRender.com, RRID:SCR_018361).

## CODE AVAILABILITY

No new software or computational algorithm was developed in this study. Custom R scripts used for compound-screen analysis, data visualization, and reproducible computational workflows are available through the verified Code Ocean compute capsule: https://codeocean.com/capsule/4710920/tree/v1. Source data required to reproduce the reported analyses are provided in the Supplementary Materials and/or public repositories as indicated.

## RESULTS

### 1. RB1-deficient osteosarcoma demonstrates high prevalence and poor neoadjuvant chemotherapy response

Analysis of the Shanghai General Hospital Osteosarcoma (SGH-OS) and Therapeutically Applicable Research to Generate Effective Treatments Osteosarcoma (TARGET-OS) cohorts showed that RB1 deficiency was present in 45% and 35% of osteosarcoma cases, respectively (**Figure 1a**). To further evaluate its clinical relevance, we retrospectively established a clinical cohort from Shanghai General Hospital with the following inclusion criteria: (1) availability of tumor whole-exome sequencing (WES) data to determine RB1 status; (2) pathological confirmation of osteosarcoma; and (3) complete pre-and post-neoadjuvant chemotherapy MRI examinations suitable for RECIST v1.1 assessment. All included patients received a standardized neoadjuvant chemotherapy regimen prior to surgery, followed by definitive surgical resection and routine postoperative chemotherapy according to institutional protocols. Radiologic evaluations were performed strictly before and after neoadjuvant chemotherapy, and histological necrosis rates were assessed on resected tumor specimens after surgery. Therefore, both RECIST-based radiologic response and pathological necrosis rate reflect response to neoadjuvant chemotherapy and are not influenced by surgical procedures or postoperative treatment.

Among the eligible patients, all RB1-non-deficient cases were included (n = 30). For comparison, 30 RB1-deficient cases were randomly selected from the remaining eligible patients using a random-number table, without knowledge of treatment response or outcome (**Figure 1b and Supplementary Table 1**).

For each patient, follow-up duration (**Figure 1b**) and radiologic response to neoadjuvant chemotherapy were recorded according to RECIST v1.1, and representative MRI scans illustrate changes in maximal tumor diameter before and after treatment in RB1-deficient and RB1-non-deficient cases (**Figure 1c**). Detailed clinical characteristics, including RB1 status, age, gender, primary tumor site, presence of recurrence or metastasis, administration of standard chemotherapy, histological necrosis rate, and post-chemotherapy disease progression status, are summarized in the clinical table (**Supplementary Table 1**). The RB1-deficient group displayed a higher proportion of progressive disease (PD) and lacked partial or complete responses, whereas RB1-non-deficient patients more frequently achieved a partial response with fewer PD events (**Figure 1d-e**). These results indicate that RB1-deficient is common in osteosarcoma and is associated with reduced sensitivity to standard neoadjuvant chemotherapy, underscoring the need to explore alternative therapeutic strategies for RB1-deficient disease.

### 2. High-throughput compound screening identifies NMS-873 as a selective inhibitor of RB1-deficient osteosarcoma

To identify potential alternative therapeutic agents for RB1-deficient osteosarcoma, we established RB1-deficient cell models (**Supplementary Figure 1a-b**) using two osteosarcoma cell lines (HOS and MG63) with either RB1 wild-type (shNC) or RB1-deficient (shRB1) status. We performed a systematic single-concentration screen of a 2,097-compound library using RB1-deficient (shRB1) and control (shNC) HOS and MG63 cells (**Figure 2a**). After four days of treatment, cell viability was measured and plate-wise normalized to DMSO controls. For each compound and each cell line, we computed the difference in normalized viability between shRB1 and shNC cells as a quantitative measure of RB1-selective sensitivity. The full raw luminescence matrix, normalized viabilities, and calculated RB1-selective differences are provided in **Supplementary Table 3**.

This analysis revealed distinct RB1-selective sensitivity distributions in both HOS and MG63 (**Figure 2b– c**). When focusing on compounds that preferentially inhibited RB1-deficient cells in both models, NMS-873(40)—a selective allosteric inhibitor of VCP—emerged as the only candidate that consistently produced strong RB1-selective suppression across both cell lines (**Figure 2d**). This reproducible pattern nominated VCP inhibition as a promising synthetic-lethal vulnerability associated with RB1-deficient status.

In validation experiments, NMS-873 treatment reduced colony formation in RB1-deficient (shRB1^#1^, shRB1^#2^) cells while showing minimal effects on RB1 wild-type (shNC) controls (**Figure 2e and Supplementary Figure 1c**). The dosing regimen for NMS-873 was selected based on IC₅₀ values determined in HOS and MG63 cells, and all concentrations used fell within the effective pharmacological range (**Supplementary Figure 1d**). These findings were further confirmed by Incucyte live-cell imaging, which demonstrated that NMS-873 treatment significantly suppressed proliferation of RB1-deficient osteosarcoma cells compared with RB1-proficient controls in RB1 knockdown HOS models (**Figure 2f**). Similar results were observed in MG63 cells (**Supplementary Figure 1e**). Three-dimensional spheroid assays further demonstrated that NMS-873 decreased the size and viability of RB1-deficient spheroids (**Figure 2g-h and Supplementary Figure 1f-g**). Notably, under DMSO treatment, RB1-deficient spheroids exhibited enhanced growth compared to RB1 wild-type spheroids, consistent with the accelerated cell cycle progression associated with RB1-deficient status. In vivo studies using a Balb/c nude mouse tibial orthotopic xenograft model (**Supplementary Figure 2a**) showed that DMSO-treated shRB1 tumors grew faster than controls, while NMS-873 treatment led to marked reduction in RB1-deficient tumor volume (**Figure 2i and Supplementary Figure 2b**) without affecting mouse body weight (**Supplementary Figure 2c**). No overt systemic toxicity was observed under this dosing schedule, as confirmed by histological examination of major organs (**Supplementary Figure 2e**). Immunohistochemical staining confirmed reduced RB1 expression in shRB1 tumors, with DMSO-treated shRB1 tumors showing significantly upregulated Proliferating Cell Nuclear Antigen (PCNA) expression (**Figure 2j-k**). These data collectively support the robustness of the cellular model system used in this study. Following NMS-873 treatment, we observed marked downregulation of PCNA accompanied by significant upregulation of the apoptosis marker cleaved Caspase-3 (c-Caspase3) (**Figure 2j-k**). Patient-derived organoid (PDO) models further validated these findings, showing potent NMS-873-mediated growth inhibition specifically in RB1-deficient osteosarcoma (**Figure 2l-m**). Immunofluorescence analysis demonstrated reduced PCNA and increased cleaved Caspase-3 in RB1-deficient PDOs after NMS-873 treatment, with minimal effects in RB1 wild-type PDOs (**Figure 2n-o**). Cell viability assays confirmed the selective inhibition of RB1-deficient PDOs by NMS-873 (**Supplementary Figure 2d**). Our comprehensive approach integrating high-throughput compound screening with validation in cellular models, animal studies, and patient-derived organoids demonstrates that the VCP inhibitor NMS-873 exerts selective and potent growth suppression in RB1-deficient osteosarcoma, highlighting its potential as an alternative targeted therapy for this molecular subtype.

### 3. Targeting VCP selectively inhibits RB1-deficient osteosarcoma

Our compound screen identified the VCP inhibitor NMS-873 as a top candidate with preferential activity against RB1-deficient osteosarcoma cells. This observation prompted further investigation of the clinical and biological relevance of VCP in this context. We constructed an osteosarcoma tissue microarray (**Supplementary Figure 3a**) and stratified patients into high/low VCP expression groups based on immunofluorescence staining (**Figure 3a and Supplementary Figure 3b-d**), while also categorizing them by disease progression status (progressive [R] vs non-progressive [nR]). Analysis revealed significantly higher VCP protein levels in the R group (**Figure 3b**). Logistic regression confirmed high VCP expression as an independent recurrence risk factor (univariate: OR=10.00, p=0.0152; multivariate: OR=7.29, p=0.0440) (**Figure 3c and Supplementary Table 4**). Kaplan-Meier analysis of TARGET-OS database showed significantly worse survival in patients with high VCP expression (**Figure 3d**). Further analysis of the TARGET-OS cohort demonstrated that high VCP mRNA levels correlated with inferior metastasis-free survival at the transcript level. Because Huvos necrosis grade is not available in the TARGET dataset and cannot be assessed from diagnostic TMA cores, multivariable models were restricted to uniformly annotated variables. Given the essential role of VCP in cellular homeostasis, we next focused on functional analyses to determine whether RB1 deficiency confers selective vulnerability to pharmacologic VCP inhibition in osteosarcoma models. Three chemically distinct VCP inhibitors (CB-5083(41), DBeQ(42), and UPCDC-30245(43)) strongly impaired clonogenic growth and proliferation in RB1-deficient HOS and MG63 cells while exerting substantially weaker effects on RB1-proficient cells (**Figure 3e and Supplementary Figure 4a**). These findings were confirmed by Incucyte live-cell imaging (**Figure 3f-h and Supplementary Figure 4b-d**). In 3D spheroid cultures (**Figure 3i and Supplementary Figure 4e-g**) and patient-derived organoids (**Figure 3j and Supplementary Figure 4h**), VCP inhibition led to marked reductions in sphere size and viability specifically in RB1-deficient models. Genetic silencing of VCP using siRNA reproduced these phenotypes, suppressing clonogenicity, proliferation, and spheroid formation in RB1-deficient cells with minimal impact on control cells (**Figure 3k-m and Supplementary Figure 5a-e**). Together, these findings demonstrate that VCP is clinically associated with aggressive disease and represents a selective therapeutic vulnerability in RB1-deficient osteosarcoma.

### 4. NMS-873 modulates VCP function through glutamine-dependent acetylation

We investigated the molecular mechanism by which NMS-873 acts on VCP. Surprisingly, NMS-873 did not affect VCP expression at either the transcriptional or protein level. We performed multiple validations at both levels: transcriptional analyses included bulk RNA-seq (**Supplementary Figure 6a–d**) and RT-PCR (**Supplementary Figure 6e**); protein-level analyses included Western-blotting (**Supplementary Figure 6f**) and immunofluorescence staining (**Supplementary Figure 6g–h**). Based on these findings, we hypothesize that NMS-873 may exert its effects by modulating the post-translational modification of VCP. Integrated analysis of Man Pan et al.’s data(38) indicated potentially binds to the VCP D2 domain, affecting acetylation at K512, K614, and K615 residues (**Supplementary Figure 7a**). Further examination of C Mori-Konya et al.’s data(39) confirmed VCP acetylation at these sites (**Figure 4a**), validated by GPS acetylation prediction software (**Supplementary Table 13**). AlphaFold3 molecular docking predicted acetyl-CoA binding at K615 (**Figure 4b**), with NMS-873 potentially interfering with this interaction (**Figure 4c-d**). GSVA analysis showed a positive correlation between acetyl-CoA biosynthesis and VCP expression in SGH-OS cohort (R=0.55, p<0.05; **Figure 4e**). To determine whether NMS-873 treatment or RB1 deficiency induces changes in histone acetylation, we examined two representative histone acetylation marks, H3K9ac and H3K27ac, by Western-blotting. No appreciable differences were observed across treatment or RB1 status (**Supplementary Figure 6i**). Co-immunoprecipitation using a pan–acetyl-lysine antibody confirmed that VCP undergoes lysine acetylation in osteosarcoma cells (**Figure 4f**). To systematically define the functional contribution of individual acetylation sites within the VCP D2 domain, we generated a panel of site-specific mutants, including K512R, K615R, an acetylation-mimetic K615Q mutant, and a combined K512R/K615R (2KR) mutant. Immunoblotting revealed that substitution of K615 with arginine (K615R) resulted in a pronounced reduction of VCP acetylation, whereas mutation of K512 alone led to only a partial decrease, and the 2KR mutant further diminished residual acetylation signals (**Figure 4g**). These results indicate that K615 represents the dominant acetylation site on VCP, with additional minor contributions from other lysine residues within the D2 domain. We next assessed whether differential acetylation states at these residues influenced cellular sensitivity to VCP inhibition. In clonogenic assays, cells expressing the acetylation-mimetic K615Q mutant exhibited reduced sensitivity to NMS-873, whereas acetylation-deficient mutants, particularly K615R and the 2KR mutant, displayed enhanced growth suppression under identical treatment conditions (**Figure 4h**). Consistent with the in vitro findings, NMS-873 significantly inhibited tumor growth only in xenografts expressing the acetylation-deficient K615R mutant, whereas xenografts expressing K615Q or K512R showed no detectable response. Moreover, under NMS-873 treatment, K615R tumors exhibited significantly greater growth suppression than K615Q tumors (**Figure 4i**). Collectively, these results establish K615 as the principal acetylation site regulating VCP function and demonstrate that modulation of VCP acetylation status at this residue directly influences cellular and tumor responses to VCP inhibition.

**Figure 4.**
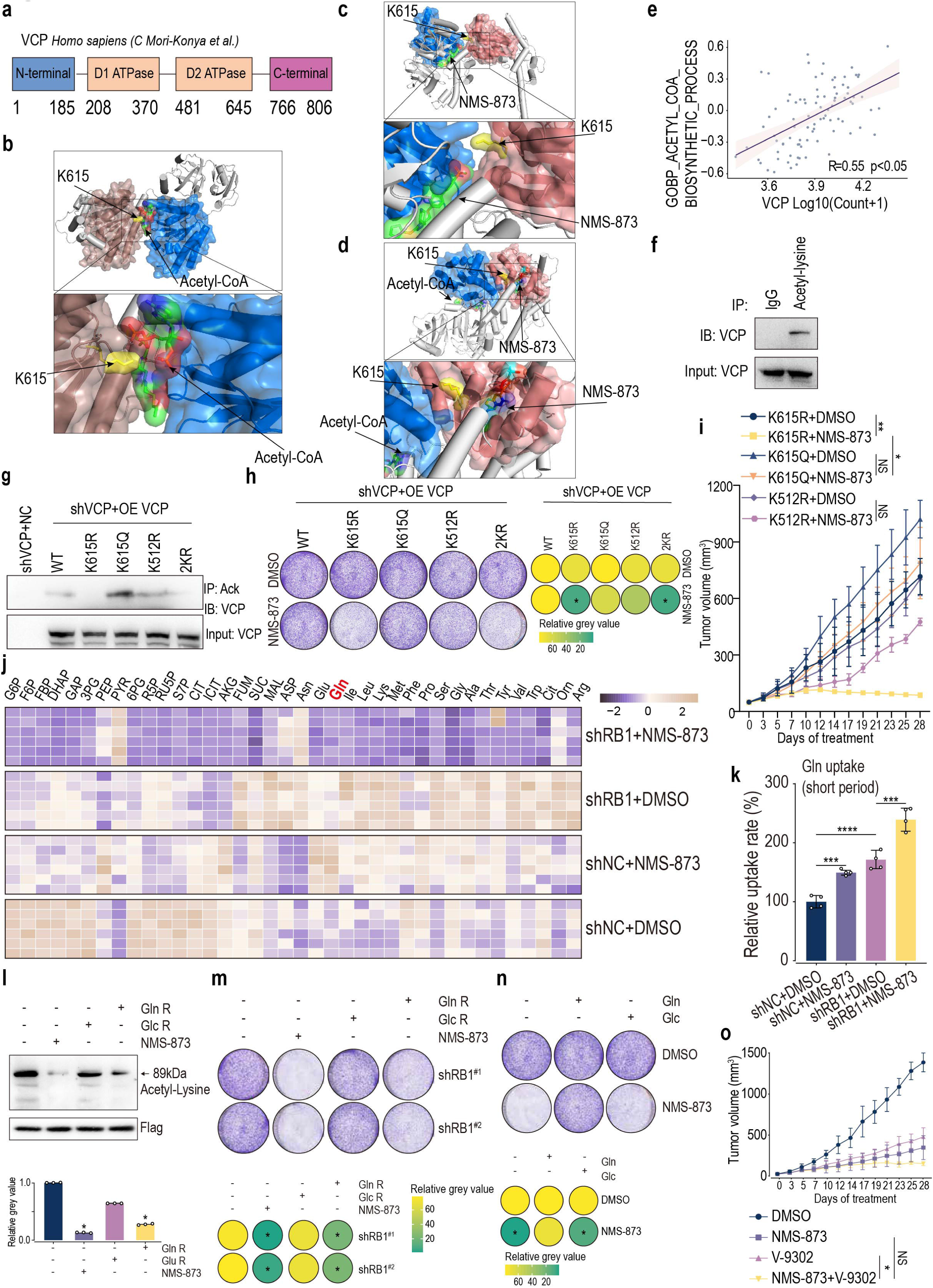
NMS-873 modulates VCP function through glutamine-dependent acetylation. **(a)** Schematic representation of the VCP protein domains. **(b)** Structural prediction using AlphaFold3 showing the interaction between acetyl-CoA and VCP at the K615 residue. The D1 domain is shown in blue, and the D2 domain in brown. **(c)** Structural prediction of the interaction between NMS-873 and VCP at K615. **(d)** Predicted model showing the spatial relationship between acetyl-CoA, NMS-873, and K615 on VCP. **(e)** Correlation analysis between the GSVA score of the acetyl-CoA biosynthetic process and VCP expression in the SGH-OS cohort (R = 0.55, p < 0.05). **(f)** Co-immunoprecipitation showing VCP acetylation in HOS osteosarcoma cells. VCP was immunoprecipitated and immunoblotted with an anti–acetyl-lysine antibody; input VCP is shown. **(g)** Acetylation levels of WT VCP and indicated lysine mutants (K615R, K615Q, K512R, and 2KR) re-expressed in a VCP-silenced background in HOS osteosarcoma cells, assessed by IP with anti–acetyl-lysine followed by IB for VCP. 2KR indicates K615R and K512R. **(h)** Clonogenic assays performed in HOS cells expressing shVCP and reconstituted with WT VCP or the indicated VCP mutants (K615R, K615Q, K512R, and 2KR), showing responses to NMS-873 (2μM) or DMSO. Data represent three biological replicates; * p < 0.05; two-way ANOVA. **(i)** Tumor growth curves of HOS-derived subcutaneous xenografts expressing the indicated VCP mutants (K615R, K615Q, or K512R). Tumor-bearing mice were treated with either DMSO or NMS-873 (20 mg/kg, intraperitoneally). Tumor growth was compared between DMSO-and NMS-873-treated groups within each mutant background, as well as between different mutant groups under NMS-873 treatment. Data represent mean ± SEM; n = 5 mice per group. Statistical analysis was performed using two-way ANOVA. **(j)** Heatmap summarizing untargeted metabolomic profiling (e.g., amino-acid-related metabolites) in HOS cells with RB1 status and NMS-873/DMSO treatment as indicated. **(k)** Glutamine uptake rate in HOS cells after NMS-873 treatment. *** indicates p < 0.001, **** indicates p < 0.0001; biological replicates: N = 4, two-way ANOVA. **(l)** Acetylation levels of VCP in HOS osteosarcoma cells under glutamine restriction (Gln R), glucose restriction (Glc R), or NMS-873 treatment. * indicates statistically significant differences with over 50% variation; biological replicates: N = 3, two-way ANOVA. **(m)** Clonogenic assays assessing the effects of nutrient restriction (Gln R or Glc R) and/or NMS-873 in RB1-deficient HOS cells, with quantification. * indicates significant difference with >50% change; two-way ANOVA. **(n)** Rescue Clonogenic assays and quantification of HOS cells under glucose or glutamine repletion following NMS-873 treatment. * indicates significant difference with >50% change; two-way ANOVA. **(o)** Tumor volume growth curves of HOS-derived RB1 wild-type (shNC) and RB1-deficient (shRB1) osteosarcoma mouse subcutaneous xenograft models treated with NMS-873, V-9302, or the combination. * indicates p < 0.05; NS indicates not significant, two-way ANOVA.

Since acetylation depends on acetyl-CoA derived from glucose or glutamine, untargeted metabolomics identified glutamine metabolism as significantly altered by NMS-873 in osteosarcoma cells (**Figure 4j**). Glutamine uptake assays revealed marked differences in osteosarcoma models (**Figure 4k, Supplementary Figure 8b and Supplementary Table 5**), with GSVA showing a positive correlation between glutamine metabolism and VCP expression in SGH-OS (R=0.55, p<0.05; **Supplementary Figure 8a and Table 5**). Only NMS-873 treatment and glutamine restriction significantly affected VCP acetylation (**Figure 4l**) and inhibited RB1-deficient osteosarcoma cell growth (**Figure 4m and Supplementary Figure 8c**), while glucose restriction showed no effect. Glutamine supplementation partially reversed NMS-873’s effects, unlike glucose (**Figure 4n and Supplementary Figure 8d**). In vivo (**Supplementary Figure 9a**), both NMS-873 monotherapy and combination with the glutamine metabolism inhibitor V-9302 significantly suppressed RB1-deficient tumor growth (**Figure 4o**), with immunohistochemistry showing reduced PCNA, enhanced c-Caspase3, and limited acetylation in combined treatment (**Supplementary Figure 9b**). These findings support a model in which NMS-873 suppresses VCP acetylation within a glutamine-regulated metabolic context, exerting potent anti-tumor effects in RB1-deficient osteosarcoma through a glutamine-dependent regulatory mechanism linking metabolism to VCP function.

### 5. RB1-deficient osteosarcoma exhibits protein folding stress

Given the heterogeneity of bone tumors, we found that analyses at the genomic level and bulk RNA-seq could not sufficiently explain the biological behavior of RB1-deficient osteosarcoma; therefore, we turned to analyzing single-cell data from osteosarcoma. We analyzed single-cell transcriptomic data from tumor samples(34). The data was partitioned into 11 cellular subgroups(n=77,998), including osteoclasts, myeloid cells, endothelial cells, lymphocytes, osteoblasts, chondroblastic cells, fibroblast subsets, and mesenchymal stem cells (**Supplementary Figure 10a**), including: (1) Osteoclast, highly expressing CTSK and MMP9; (2) Myeloid, highly expressing HLA-DRA and C1QA; (3) Endothelial, highly expressing PECAM1 and VWF; (4) Lymphocyte, highly expressing CD3D and NKG7; (5) Osteoblast, highly expressing ALPL and RUNX2; (6) Chondroblastic, highly expressing COL2A1, SOX9 and ACAN; (7) Fibroblast_01, highly expressing TAGLN, ACTA2 and RGS5; (8) Fibroblast_02, highly expressing DCN, and COL1A1; (9) Fibroblast_03, highly expressing DCN, C1S and COL1A1; (10) MSC_01, highly expressing MKI67, TOP2A and CCNB1; (11) MSC_02, highly expressing PCNA (**Supplementary Figure 10b**).

We focused specifically on the mesenchymal lineage subpopulations, including osteoblasts, chondroblastic cells, three fibroblast subpopulations (Fibroblast_01, Fibroblast_02, and Fibroblast_03), as well as proliferative mesenchymal stem cell (MSC) subpopulations (MSC_01 and MSC_02) (**Supplementary Figure 10c**). We assessed the expression of key proliferation markers within these cell groups. UMAP visualization showed that the expression levels of MKI67 and PCNA were significantly elevated in the proliferative MSC subpopulations (**Supplementary Figure 10d**). We calculated the copy number variation (CNV) scores within the mesenchymal lineage populations (**Supplementary Figure 10e**). Based on RB1 CNV scores, we grouped the cells showing both high PCNA/MKI67 expression and high CNV scores. Differential pathway analysis revealed significant upregulation of 112 pathways in RB1-deficient tumor cells compared with RB1-non-deficient tumor cells (**Supplementary Table 6 and Figure 5a**). Through integrated analysis of MG63 RNA-seq (shRB1 vs. shNC, **Supplementary Table 7**), single-cell RNA-seq, and VCP-related pathway data from SGH-OS (**Supplementary Table 8**) and TARGET-OS (**Supplementary Table 9**) cohorts, we identified 38 consistently upregulated pathways across all four datasets (**Figure 5b**). These pathways primarily involved eight major biological categories (**Supplementary Figure 10f**), including: endoplasmic reticulum stress and protein folding, mitochondrial gene expression and translation, mitochondrial structure and transport, ribosome biogenesis and protein translation, tRNA processing and modification, other RNA processing and modification, metabolism, and small molecule biosynthesis. Among these, endoplasmic reticulum stress and protein folding pathways represented the highest proportion, with overlapping pathways including: cellular response to topologically incorrect proteins, cellular response to unfolded proteins, endoplasmic reticulum unfolded protein response, establishment of protein localization to organelles, intrinsic apoptotic signaling pathway, protein folding, regulation of endoplasmic reticulum unfolded protein response, and regulation of intrinsic apoptotic signaling pathway (**Figure 5c**). These findings suggest that RB1-deficient osteosarcoma exhibits significant protein folding stress.

**Figure 5.**
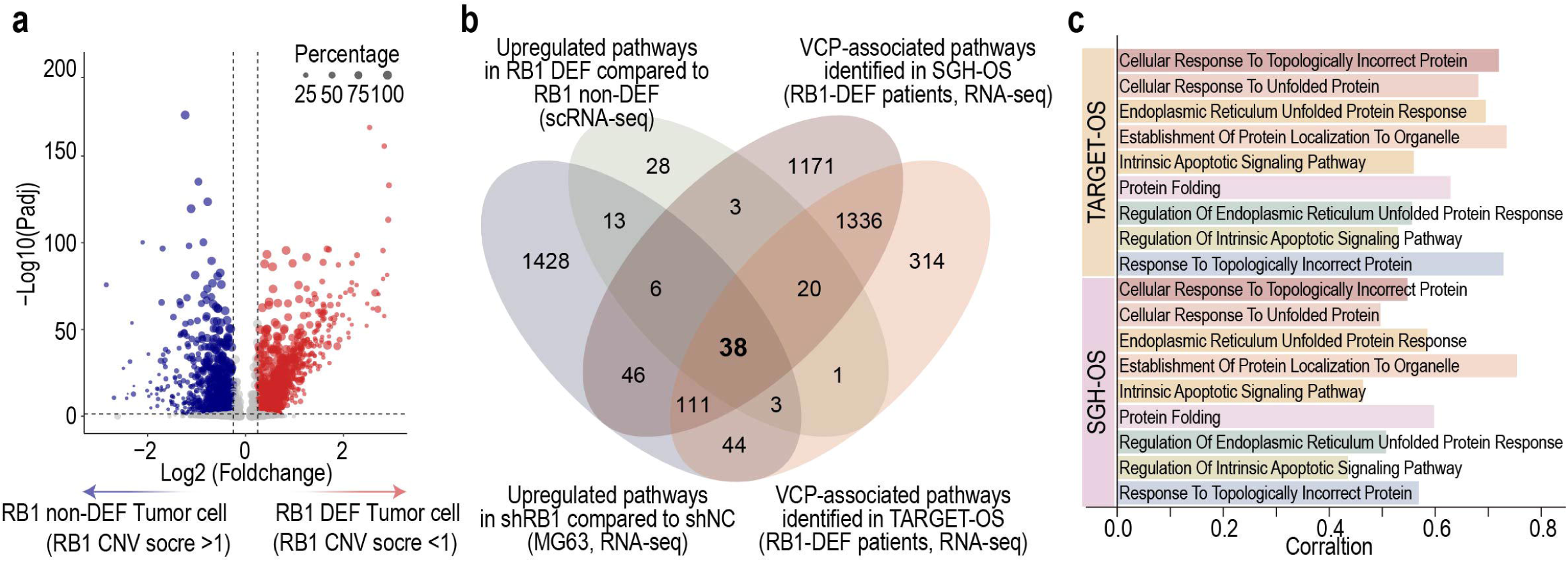
RB1-deficient osteosarcoma exhibits protein folding stress. **(a)** Volcano plot showing differentially expressed pathways between RB1-deficient and non-deficient tumor cells, stratified based on RB1 CNV score. **(b)** Venn diagram summarizing the overlap of upregulated pathways from four datasets: (1) shRB1 vs. shNC in MG63 RNA-seq, (2) RB1-DEF vs. non-DEF in single-cell RNA-seq, (3) VCP-associated pathways in RB1-deficient patients from the SGH-OS cohort, and (4) VCP-associated pathways in RB1-deficient patients from the TARGET-OS cohort. **(c)** Correlation of VCP-associated pathways in RB1-deficient patients from the SGH-OS and TARGET-OS RNA-seq datasets. Pathways include: Cellular response to topologically incorrect protein, Cellular response to unfolded protein, Endoplasmic reticulum unfolded protein response, Establishment of protein localization to organelle, Intrinsic apoptotic signaling pathway, Protein folding, Regulation of endoplasmic reticulum unfolded protein response, Regulation of intrinsic apoptotic signaling pathway, Response to topologically incorrect protein.

To further characterize the cellular communication landscape in osteosarcoma, we performed CellChat ligand–receptor analysis across identified tumor cell populations. Examination of signaling programs associated with cytokine and stress-related communication showed that RB1-deficient tumors exhibited higher communication probabilities in MIF–CD74/CD44 and JAG–NOTCH signaling across multiple mesenchymal lineages (**Supplementary Figure 11a-b**). Expanded CellChat profiling also demonstrated differences in the global organization of ligand–receptor signaling between RB1-deficient and non-deficient tumors. Outgoing and incoming signaling heatmaps revealed broader and more frequent signaling interactions in proliferative MSC and fibroblast subsets in the RB1-deficient group (**Supplementary Figure 11c-d**). Network analyses further showed that the RB1-deficient group had a greater number of interactions and greater interaction weights across mesenchymal and immune populations (**Supplementary Figure 11e-f**). Taken together, these analyses show that RB1-deficient osteosarcoma displays increased ligand–receptor communication activity both within mesenchymal populations and between mesenchymal and immune cell types, compared with RB1-non-deficient tumors.

### 6. Inhibition of VCP acetylation induces endoplasmic reticulum stress in RB1-deficient osteosarcoma cells

Gene-set variation analysis of the RNA-seq datasets from HOS and MG63 cells showed enrichment of pathways related to regulation of translation initiation in response to endoplasmic reticulum (ER) stress following NMS-873 treatment (**Figure 6a and Supplementary Table 10-11**). Consistent with these transcriptomic changes, immunofluorescence staining revealed increased aggresome accumulation in HOS cells after NMS-873 exposure (**Figure 6b–c**), with similar results observed in MG63 cells (**Supplementary Figure 12a**). Apoptosis assays demonstrated that RB1-deficient cells exhibited higher Caspase-3/7 activity after treatment with classical ER stress inducers (thapsigargin or tunicamycin) as well as with NMS-873 (**Figure 6d–e and Supplementary Figure 12b**). Colony-formation assays further showed that these ER-stress–related agents reduced clonogenic survival of RB1-deficient osteosarcoma cells (**Figure 6f and Supplementary Figure 12c**). In rescue experiments, treatment with RH01687 or RH01386 partially restored the colony-forming ability of both HOS and MG63 cells under NMS-873 exposure (**Figure 6g**). RH01687 and RH01386 are small-molecule compounds reported to alleviate endoplasmic reticulum stress by improving protein homeostasis and reducing unfolded protein accumulation(44). RT-qPCR analysis also showed elevated mRNA expression of these ER-stress–related genes after NMS-873 treatment (**Figure 6h and Supplementary Figure 12d**). Western-blotting showed increased protein levels of multiple ER-stress pathway components—including IRE1, XBP1, PERK, EIF2α, CHOP, ATF6, and HSPA5—in RB1-deficient cells treated with NMS-873 compared with vehicle controls (**Figure 6i and Supplementary Figure 12e**). Immunohistochemical staining of tumors derived from the tibial orthotopic xenograft model demonstrated elevated expression of multiple ER stress– associated proteins in RB1-deficient osteosarcoma following NMS-873 treatment, consistent with the ER stress activation observed in vitro (**Figure 6j**). In glutamine-restriction assays, removal of glutamine enhanced ER-stress marker expression, whereas glutamine supplementation reduced the induction of several ER-stress genes under NMS-873 treatment (**Figure 6k**). Taken together, multiple orthogonal assays—including transcriptomic enrichment, aggresome staining, apoptosis activation, colony formation, Western-blotting, and qPCR—consistently show that NMS-873 treatment is accompanied by activation of ER-stress–related signaling pathways in RB1-deficient osteosarcoma cells.

**Figure 6.**
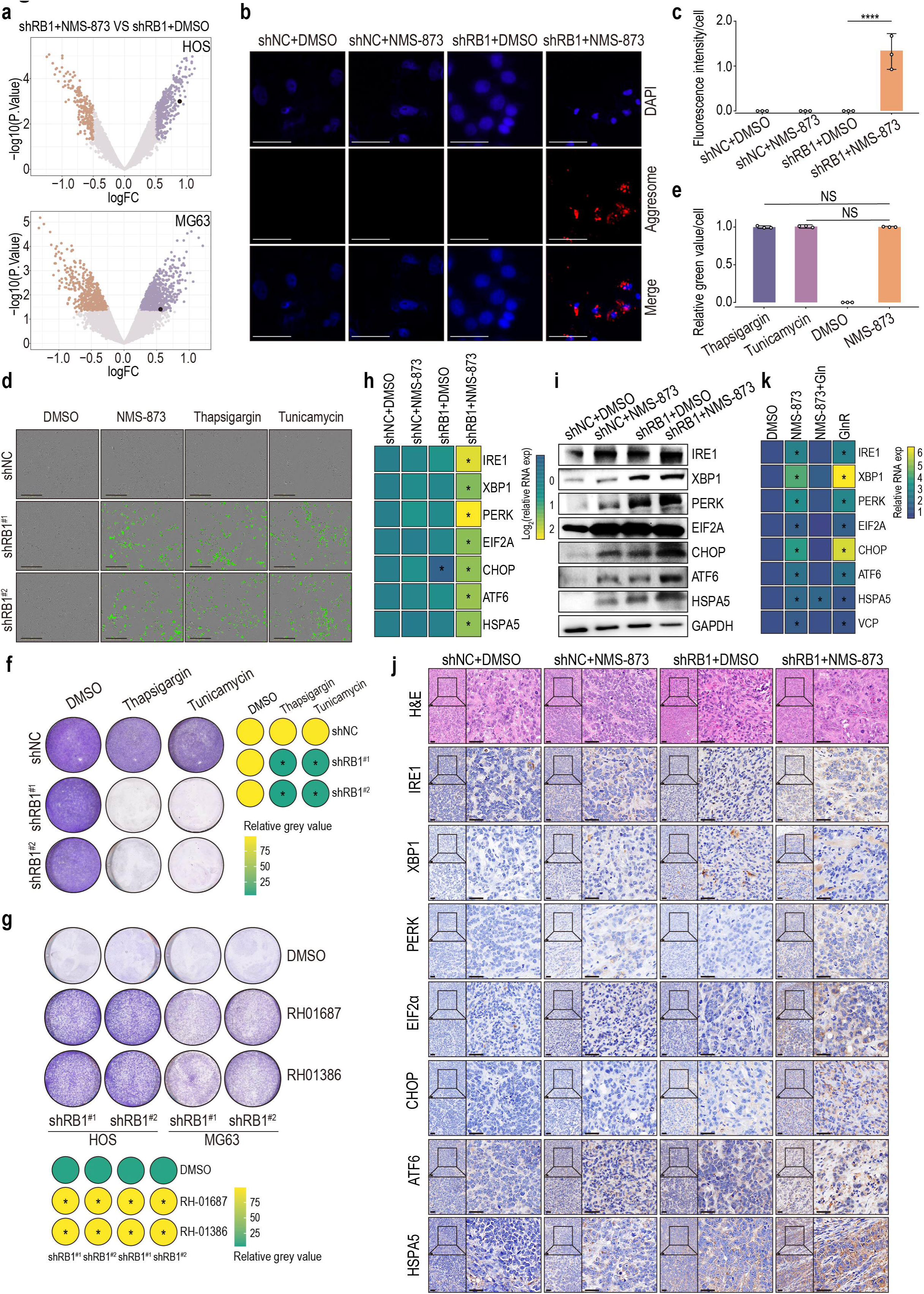
Inhibition of VCP acetylation induces endoplasmic reticulum stress in RB1-deficient osteosarcoma cells. **(a)** GSVA analysis of HOS and MG63 RNA-seq data under NMS-873 treatment. Each point represents a GSVA pathway score. The black dot indicates the pathway “regulation of translation initiation in response to endoplasmic reticulum (ER) stress”. **(b)** Immunofluorescence staining of aggresomes in HOS cells treated with NMS-873. Scale bar = 100μm. **(c)** Quantification of aggresome-positive cells in HOS cells treated with NMS-873. Technical replicates: N = 3. **** indicates p < 0.0001; analyzed by two-way ANOVA. **(d)** Caspase-3/7 activation in HOS cells treated with thapsigargin or tunicamycin. Scale bar = 400μm. Representative images are shown; quantification is shown in (**e**). **(e)** Quantification of Caspase-3/7 fluorescence in HOS cells after thapsigargin or tunicamycin treatment. Technical replicates: N = 3. NS indicates not significant (p > 0.05); analyzed by two-way ANOVA. **(f)** Clonogenic assays showing the effect of thapsigargin and tunicamycin on HOS cell viability. Biological replicates: N = 3. * indicates p < 0.05; two-way ANOVA. **(g)** Clonogenic assays showing the effects of RH01687 and RH01386 under NMS-873 treatment in HOS and MG63 cells. Biological replicates: N = 3. * indicates p < 0.05; two-way ANOVA. **(h)** RT-qPCR analysis of mRNA levels for IRE1, XBP1, PERK, EIF2α, CHOP, ATF6, and HSPA5 in HOS cells after NMS-873 treatment. Biological replicates: N = 3. * indicates p < 0.05; two-way ANOVA. **(i)** Western-blot analysis of ER stress-related proteins (IRE1, XBP1, PERK, EIF2α, CHOP, ATF6, and HSPA5) in HOS cells after NMS-873 treatment. **(j)** Immunohistochemical staining of ER stress–related markers (IRE1, XBP1, PERK, EIF2α, CHOP, ATF6, and HSPA5) in tumor tissues derived from the tibial orthotopic xenograft model following NMS-873 treatment. Scale bar, 50μm. **(k)** RT-qPCR analysis of ER stress-related gene expression (IRE1, XBP1, PERK, EIF2α, CHOP, ATF6, HSPA5, and VCP) in HOS cells under glutamine restriction, NMS-873 treatment, or both. Biological replicates: N = 3. * indicates p < 0.05; two-way ANOVA.

## DISCUSSION

RB1, a critical tumor suppressor, loses function due to genomic mutations, resulting in uncontrolled cell proliferation and tumorigenesis(45, 46). The accelerated cell cycle in RB1-deficient cells leads to increased protein synthesis during interphase, imposing heightened stress on the endoplasmic reticulum (ER). This mechanistic link explains the sensitivity of RB1-deficient osteosarcoma cells to NMS-873, a small-molecule inhibitor of VCP, a key regulator of ER homeostasis(40). Our findings underscore the importance of investigating ER homeostasis regulation in RB1-deficient tumor cells, which could reveal fundamental insights into their survival mechanisms and offer novel therapeutic strategies for targeting these vulnerable pathways in RB1-deficient cancers.

Our study further demonstrates that the loss of RB1 function leads to metabolic reprogramming of glutamine in tumor cells, which plays a crucial role in regulating endoplasmic reticulum (ER) homeostasis. In RB1-deficient osteosarcoma, glutamine metabolism not only provides energy but also contributes to the remodeling of epigenetic landscapes and post-translational modifications. Glutamine metabolism may contribute to the acetyl-CoA pool that supports non-histone protein acetylation, including VCP acetylation. Notably, NMS-873 binds to VCP at a region encompassing the acetylation site lysine 615 (K615). Upon NMS-873 treatment, acetylation of VCP in RB1-deficient osteosarcoma cells is reduced. Parallel experiments revealed that only glutamine restriction, and not carbohydrate restriction, could replicate this effect, suggesting a critical link between VCP function in RB1-deficient osteosarcoma and glutamine-mediated protein acetylation.

Further investigation revealed that glutamine supplementation significantly attenuated the cytotoxic effects of NMS-873 in RB1-deficient osteosarcoma cells, underscoring the functional importance of glutamine availability in this context. To assess whether these effects were accompanied by alterations in histone acetylation, we examined two representative histone acetylation marks, H3K9ac and H3K27ac. No appreciable differences were detected following NMS-873 treatment. These observations argue against a prominent role for broad histone acetylation changes under these conditions and instead support a model in which glutamine availability predominantly regulates acetylation of specific non-histone substrates, notably VCP, rather than driving generalized chromatin-level acetylation changes.

In summary, our study identifies VCP, a key protein involved in maintaining endoplasmic reticulum (ER) homeostasis(47–49), as a novel synthetic lethal target in RB1-deficient osteosarcoma, along with the small-molecule inhibitor NMS-873 that targets VCP(50). We established a link between glutamine metabolic reprogramming following RB1 loss and the synthetic lethality mechanism involving VCP. NMS-873 reduces acetylation of VCP, leading to its functional inactivation, thereby exacerbating ER stress in RB1-deficient osteosarcoma cells and inducing cell death. It is important to note that NMS-873 has been biochemically characterized as an allosteric inhibitor with strong preference for VCP/p97 over other AAA+ ATPases. In our study, several lines of evidence collectively support that the phenotypes observed are predominantly mediated through on-target VCP inhibition. Genetic suppression of VCP using siRNA closely reproduced the growth-inhibitory patterns induced by NMS-873, particularly the heightened vulnerability of RB1-deficient cells. Moreover, two structurally unrelated VCP inhibitors elicited similar RB1-selective responses, and mutation of the functionally relevant acetylation site altered drug sensitivity in a manner consistent with VCP-dependent regulation. Although off-target effects cannot be entirely excluded for any small-molecule inhibitor, the concordance among pharmacologic perturbation, genetic depletion, and mutation-based modulation strongly supports that the synthetic lethality observed here is largely attributable to VCP inhibition.

Beyond mechanistic specificity and current progress in the development of VCP inhibitors, our findings have potential implications for refining the therapeutic window of VCP-targeted therapy. RB1-deficient osteosarcoma cells consistently displayed markedly enhanced sensitivity to NMS-873, with substantially lower IC₅₀ values than RB1-proficient counterparts. This raises the possibility that clinically meaningful antitumor activity might be achievable at lower systemic exposures in RB1-deficient tumors compared with unselected populations. In principle, such a genetically stratified approach could mitigate some of the dose-limiting toxicities that hindered the development of first-generation VCP inhibitors, although dedicated pharmacokinetic and toxicology studies will be required to formally assess this potential. Together, these observations provide a biologically grounded framework for future efforts aimed at translating VCP inhibition into a therapeutically actionable strategy for RB1-deficient osteosarcoma.

Given the rarity of osteosarcoma, clinical translational research faces inherent challenges. Through a small-scale cohort, we preliminarily confirmed the value of VCP in osteosarcoma recurrence, suggesting that VCP plays a critical role not only in RB1-deficient osteosarcoma but also in the overall malignancy of osteosarcoma, where ER homeostasis stability appears to be linked to disease severity. Future studies should explore the clinical translational potential of NMS-873 in osteosarcoma treatment and address additional unresolved questions highlighted in this work.

## CONSENT FOR PUBLICATION

Not applicable.

## RESOURCE AVAILABILITY

### Lead contact

Further information and requests for resources and reagents should be directed to and will be fulfilled by the lead contact, Wei Sun.

### Materials availability

This study did not generate new unique reagents.

### Data and code availability

Data: All data generated and analyzed in this study are provided in the Supplementary Materials.

Code: This study did not result in any development of original code. Custom R scripts used for compound screen analysis, data visualization, and reproducible computational workflows are available through the verified Code Ocean compute capsule: https://codeocean.com/capsule/4710920/tree/v1

Any additional information required to reanalyze the data reported in this work paper is available from the lead contact upon request.

## ACKNOWLEDGMENTS

We would be deeply grateful to Professor Cun Wang [State Key Laboratory of Oncogenes and Related Genes, Shanghai Cancer Institute, Renji Hospital, Shanghai Jiao Tong University School of Medicine, Shanghai, China] for his valuable suggestions and feedback that greatly improved this work. We would be deeply grateful to Professor Wen Wen [Department of Laboratory Diagnosis, Third Affiliated Hospital of Naval Medical University (Second Military Medical University), Shanghai, China.], Professor Bing Li [Department of Biochemistry and Molecular Cell Biology, Shanghai Key Laboratory for Tumor Microenvironment and Inflammation, Shanghai Jiao Tong University School of Medicine], and Jianyuan Zhao [Xinhua Hospital, Shanghai Jiao Tong University School of Medicine, Shanghai, China] for their valuable suggestions and help.

This work was supported in part by the National Natural Science Foundation of China (82373177 to W. Sun, 82473469 to H.S. Wang, 82404064 to H.R. Mu, 82404063 to K.Y. Liu, 82473491 and 82172366 to L. Yang).

## AUTHOR CONTRIBUTIONS

W.S., H.S.W. and D.Q.Z. designed the research. H.R.M. and B.H.Y. wrote the manuscript. H.R.M., D.Q.Z., H.S.W., J.K.S., Z.Y.W. and K.Y.L. collected the clinical data and specimens. B.H.Y., Y.C., H.R.M, Q.Z., Y.N.T., X.H., X.Y.Y., H.Y.W. and Y.Y. performed the experiments. H.R.M., H.Y.W., B.W.Z., and Y.N.T. performed bioinformatic analyses. Z.D.C., Y.Q.H., T.Z., L.Y., and J.X. supervise the experiments. Z.Y.W. managed the reagents and expenses used in the project. All authors have read and approved the article.

---

**Supplementary Figure 1.**
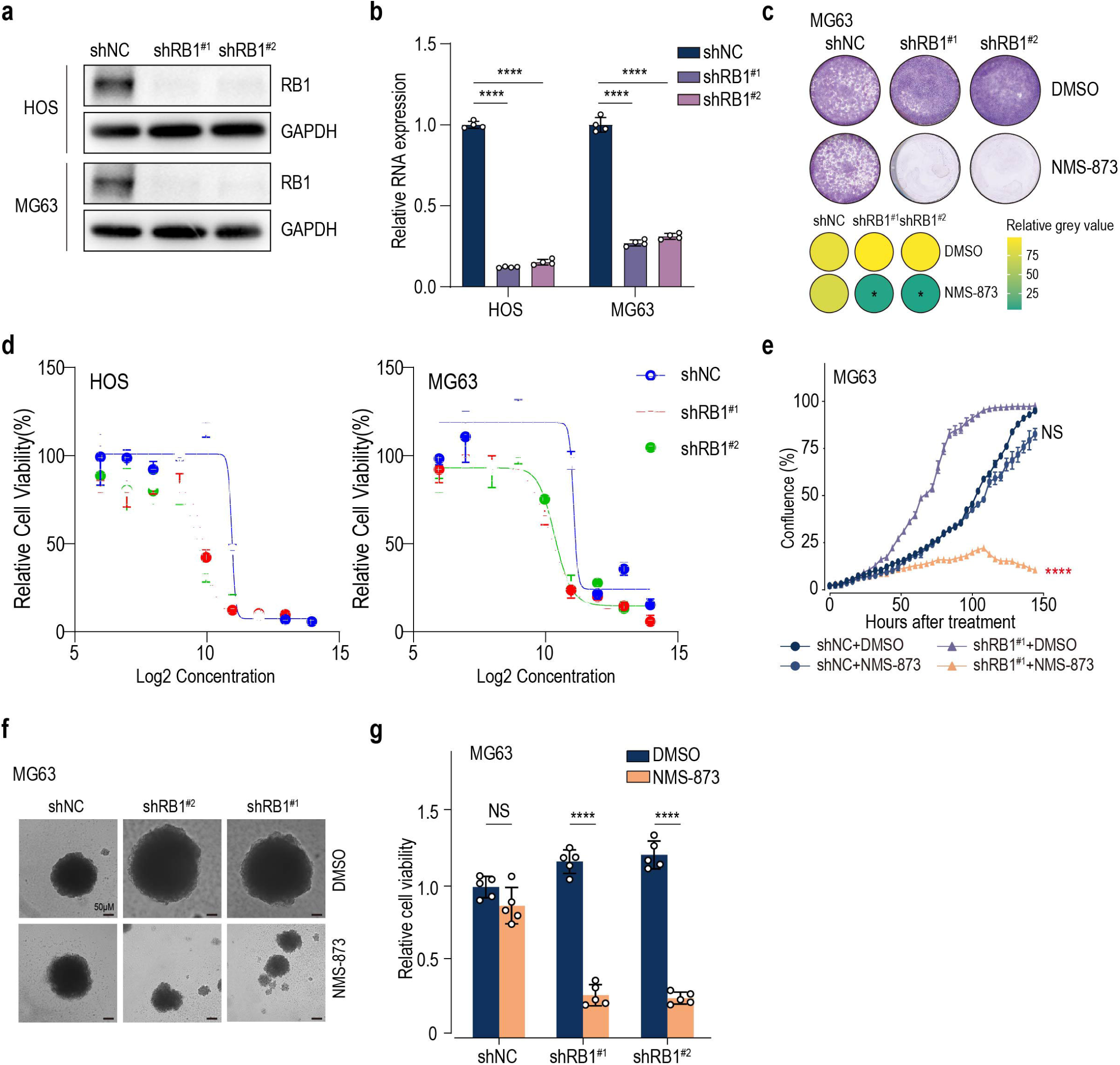

**Supplementary Figure 2.**
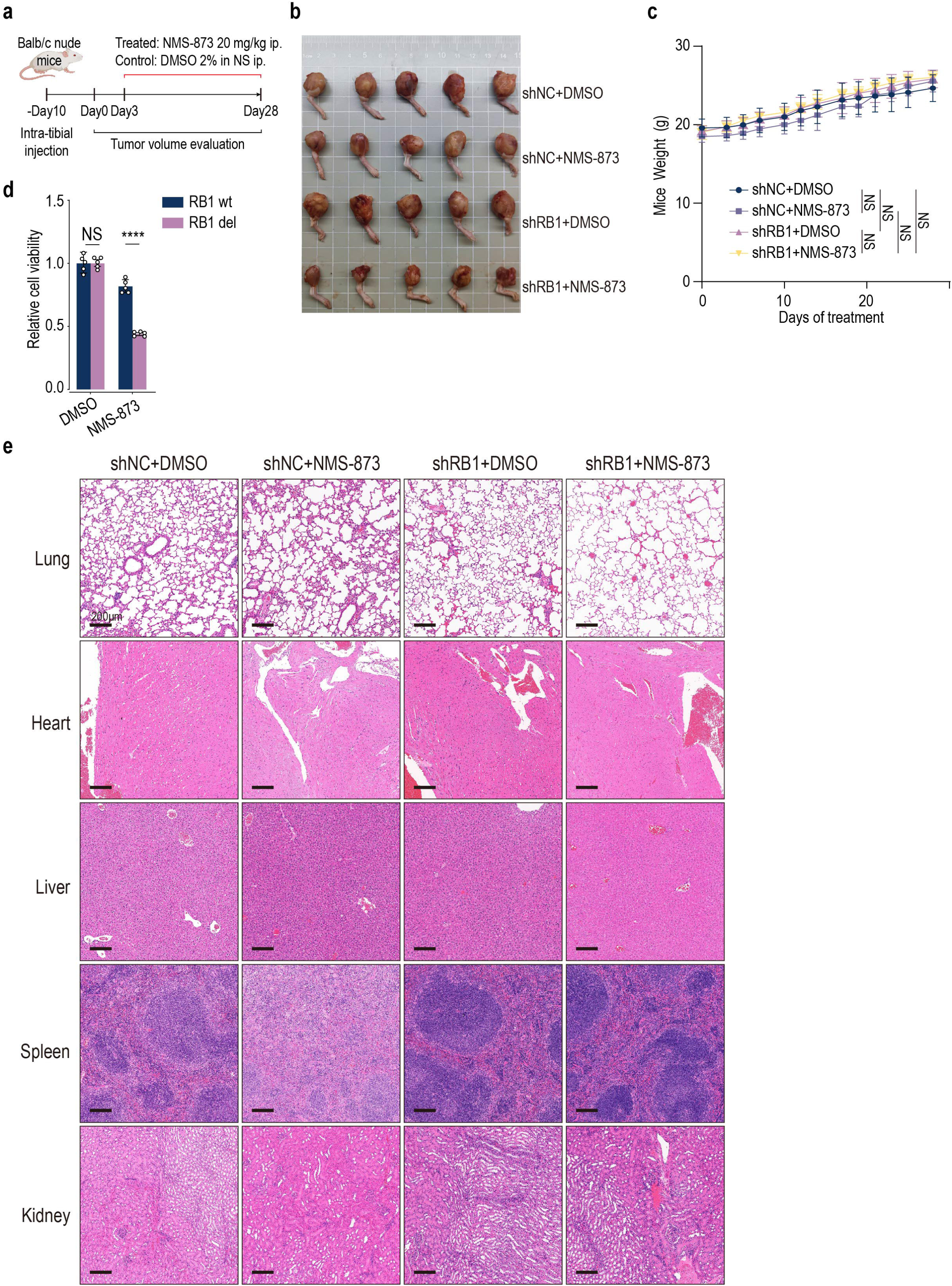

**Supplementary Figure 3.**
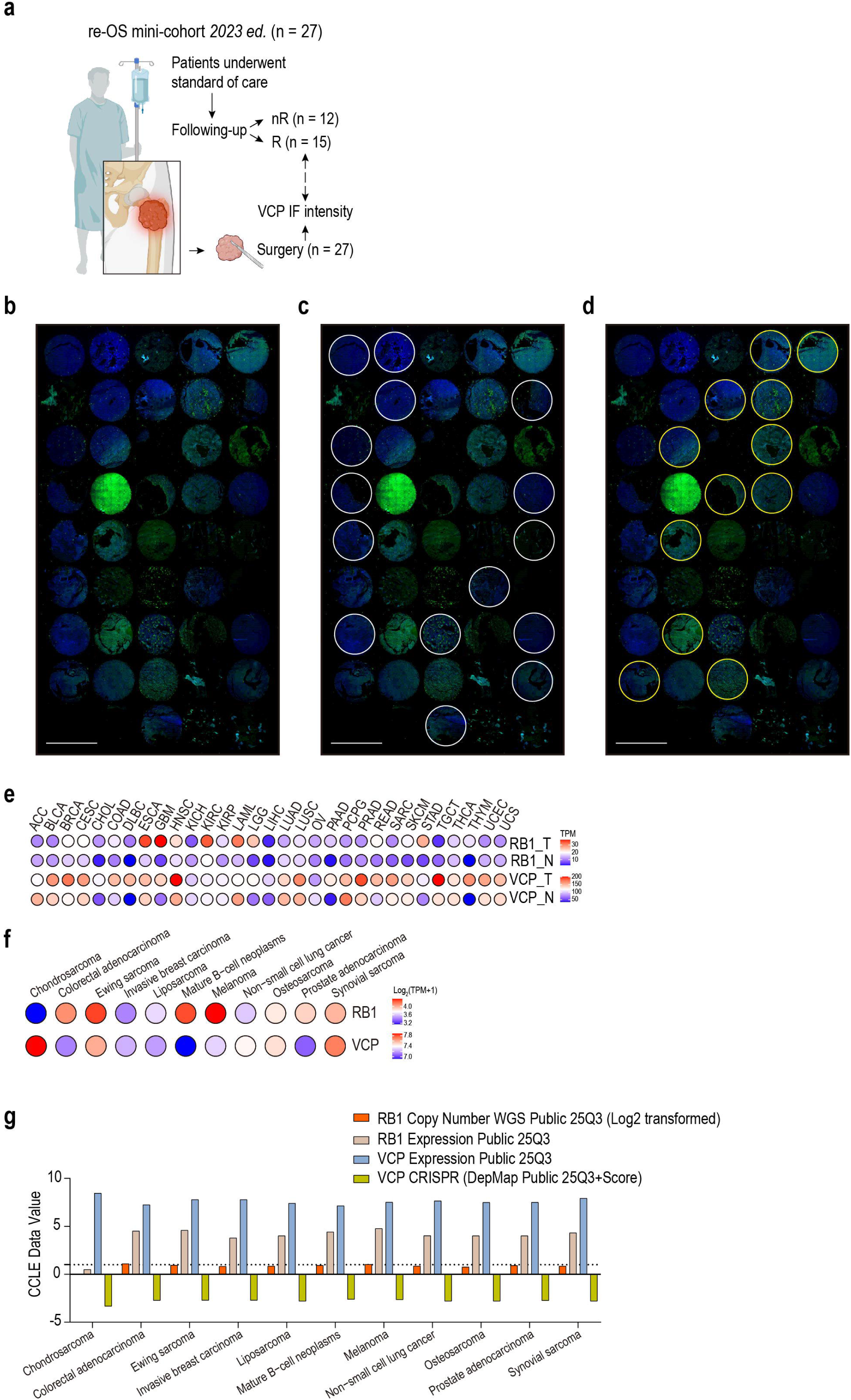

**Supplementary Figure 4.**
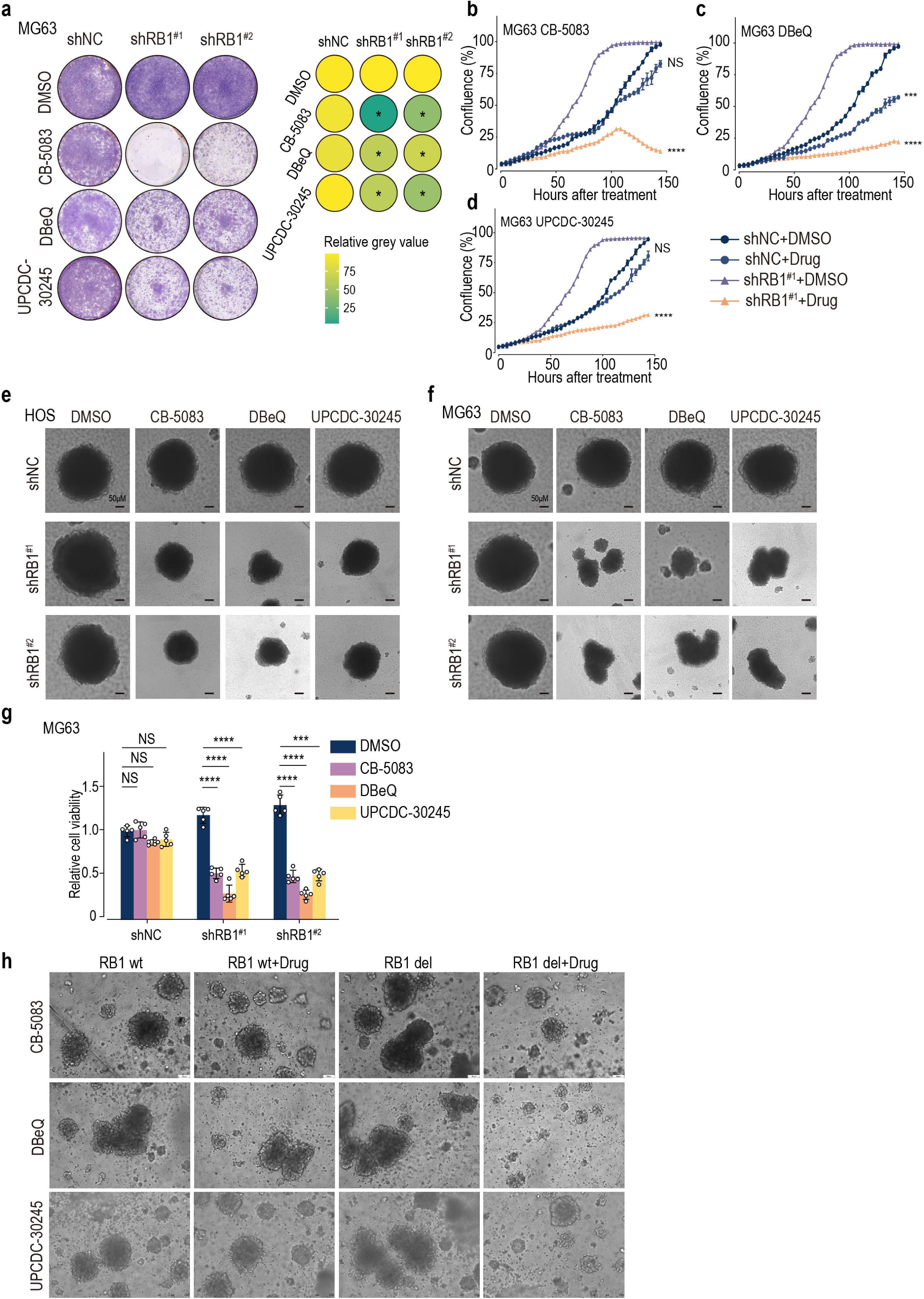

**Supplementary Figure 5.**
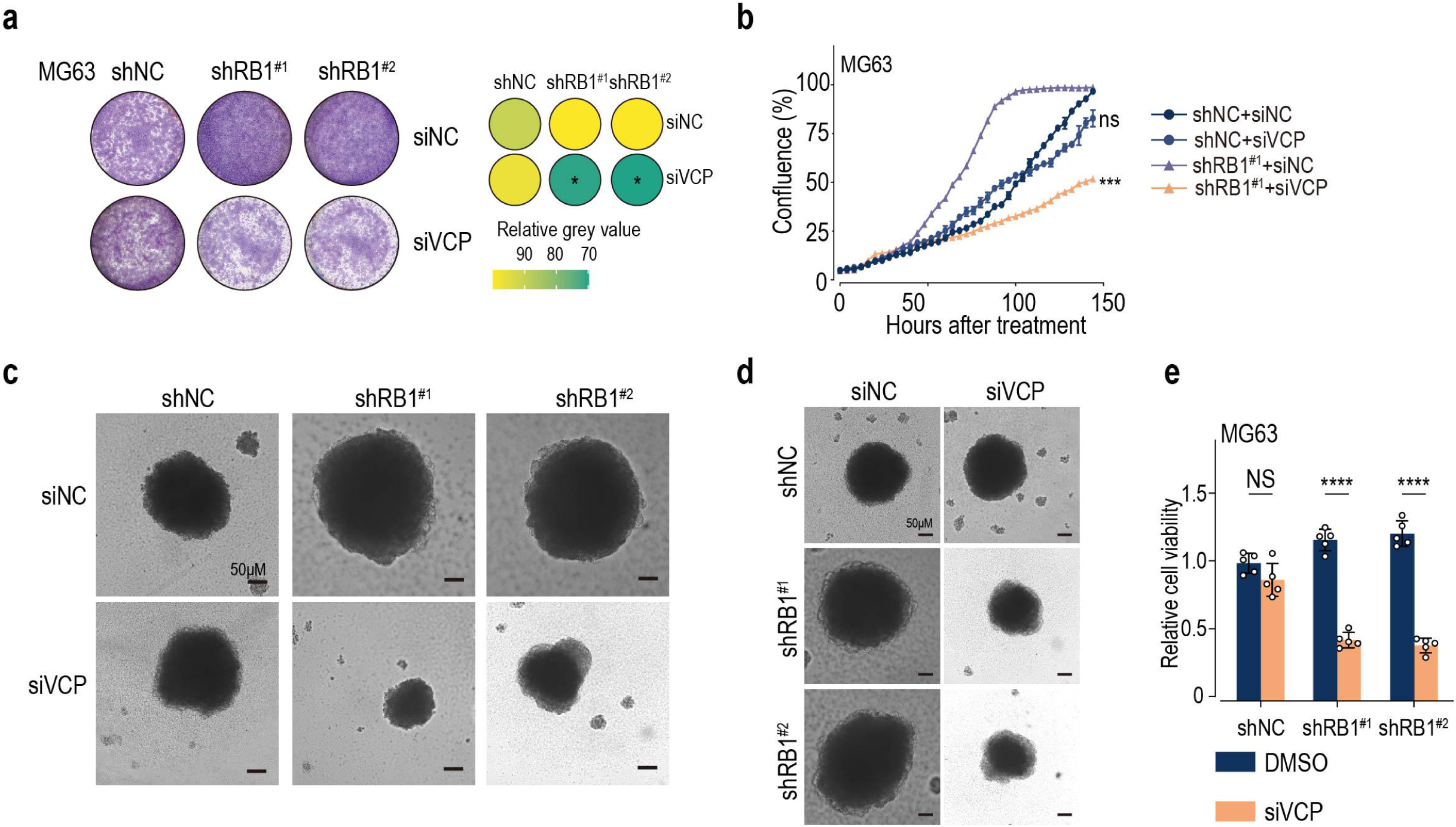

**Supplementary Figure 6.**
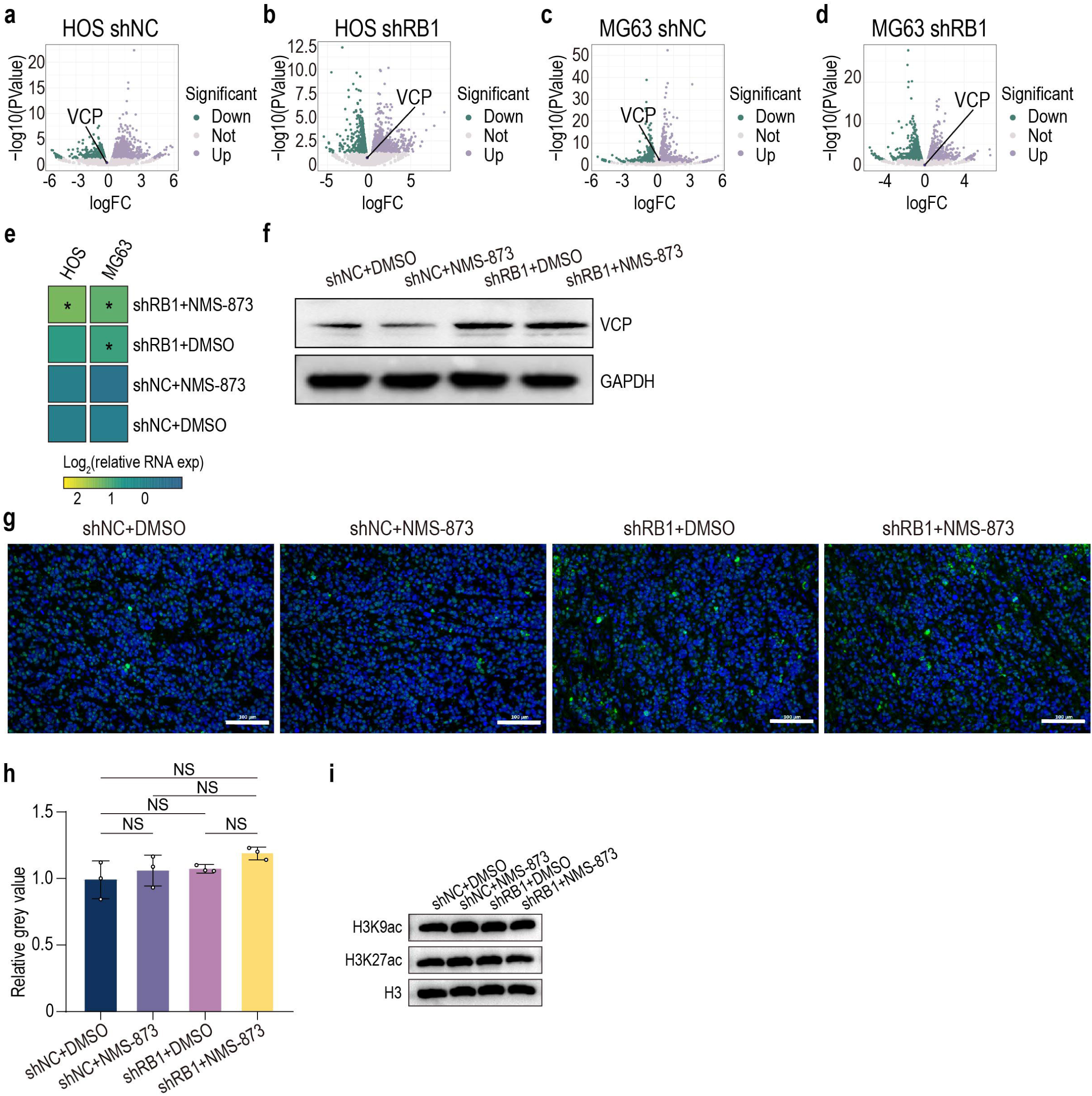

**Supplementary Figure 7.**
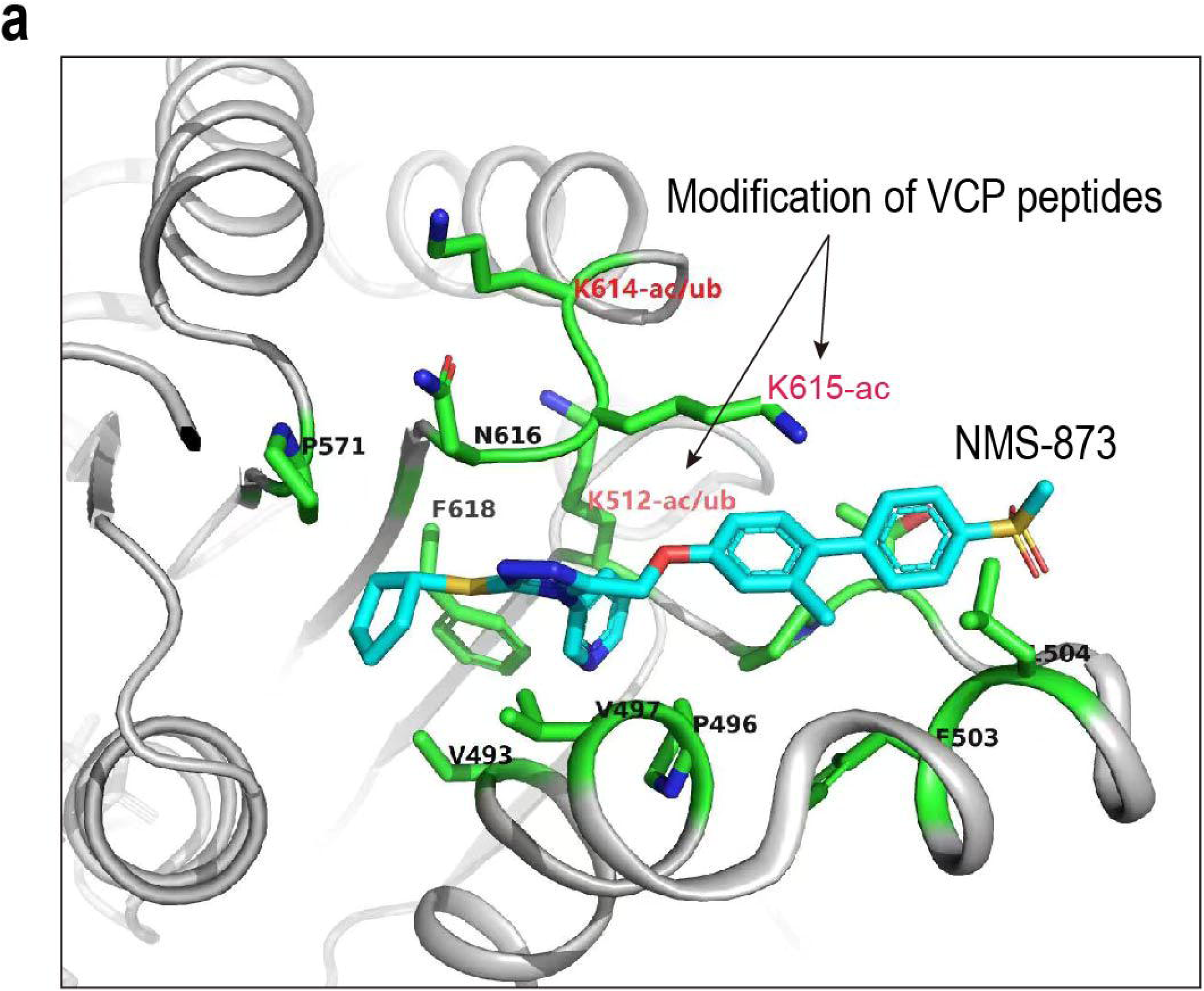

**Supplementary Figure 8.**
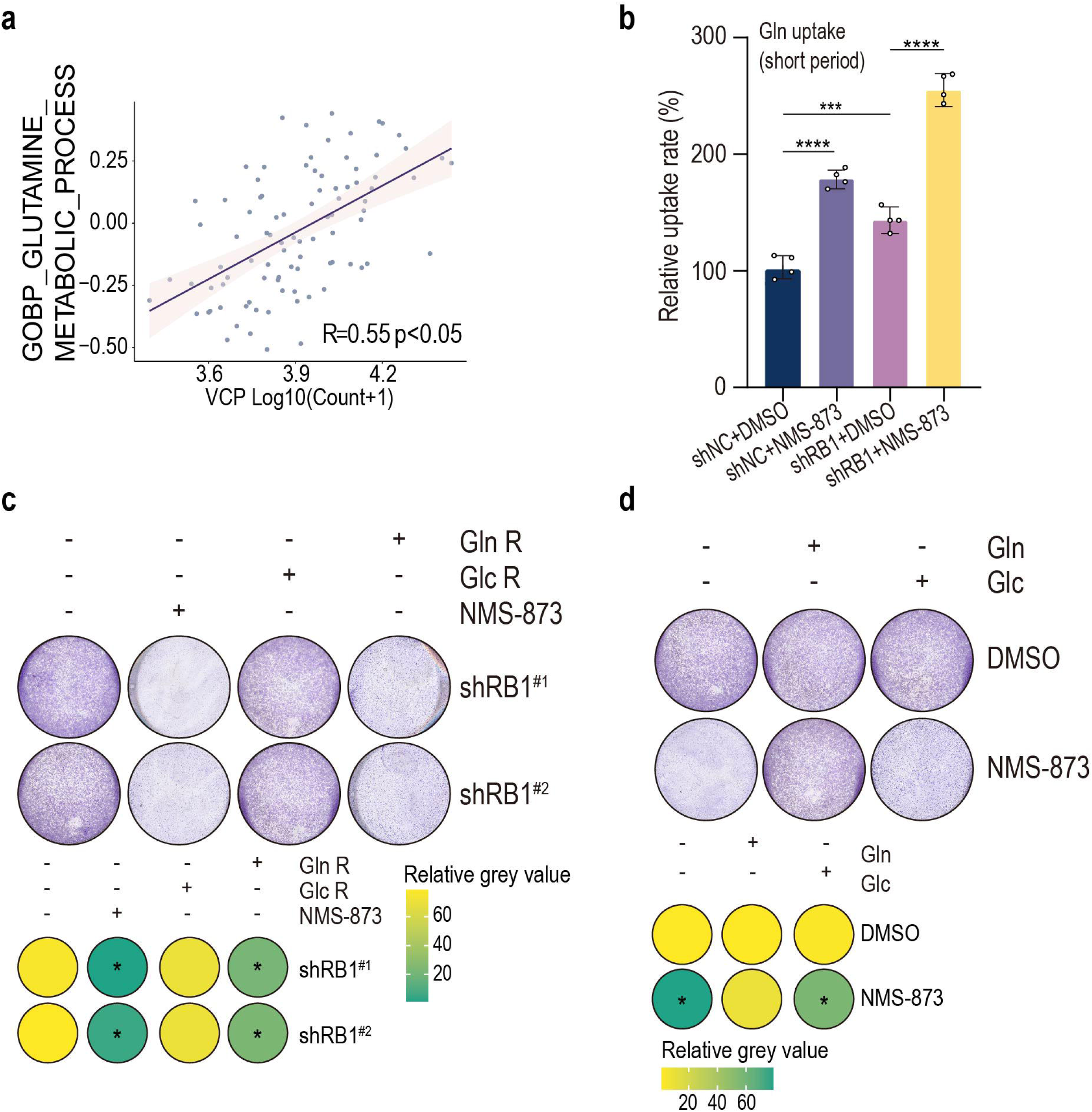

**Supplementary Figure 9.**
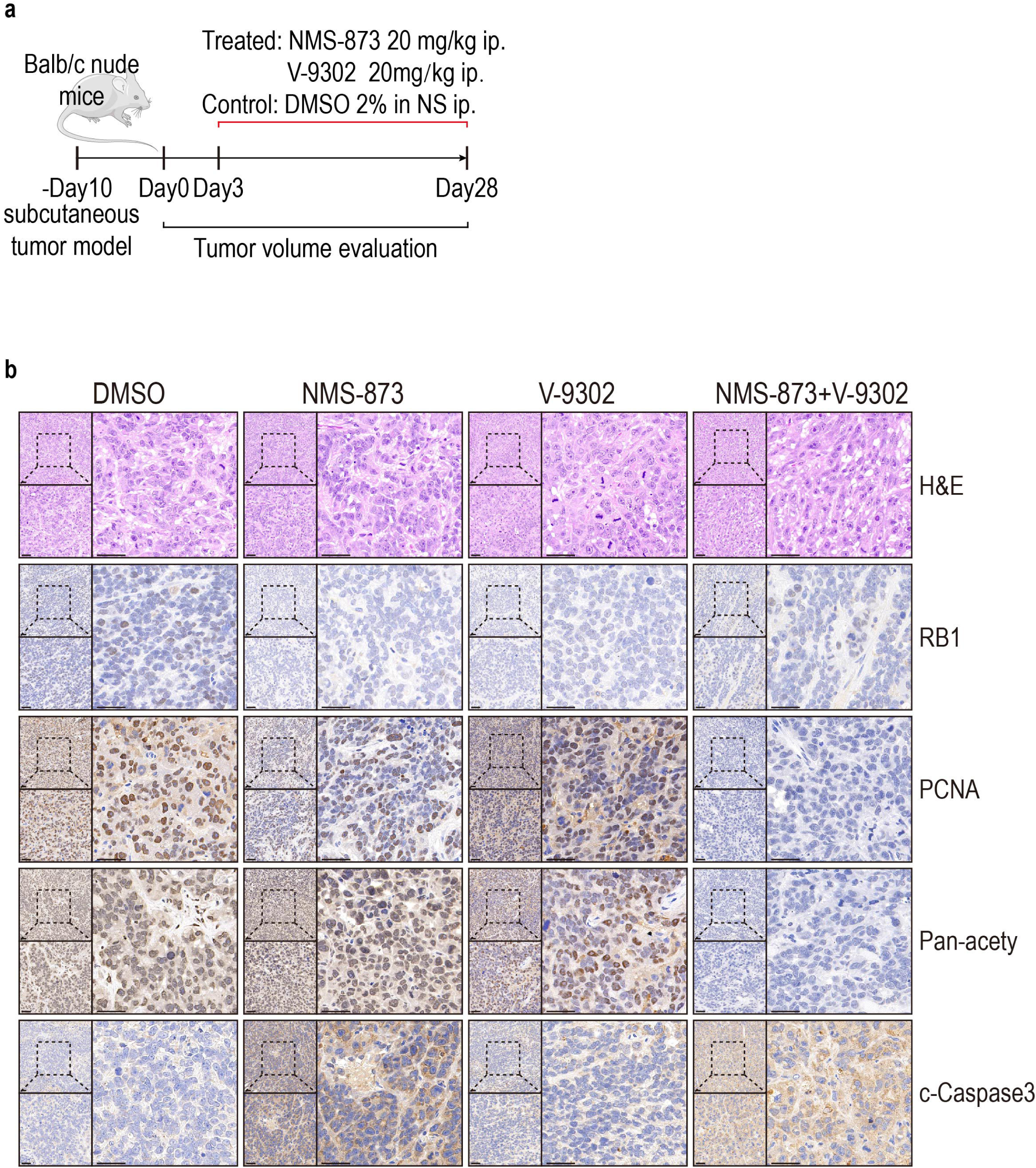

**Supplementary Figure 10.**
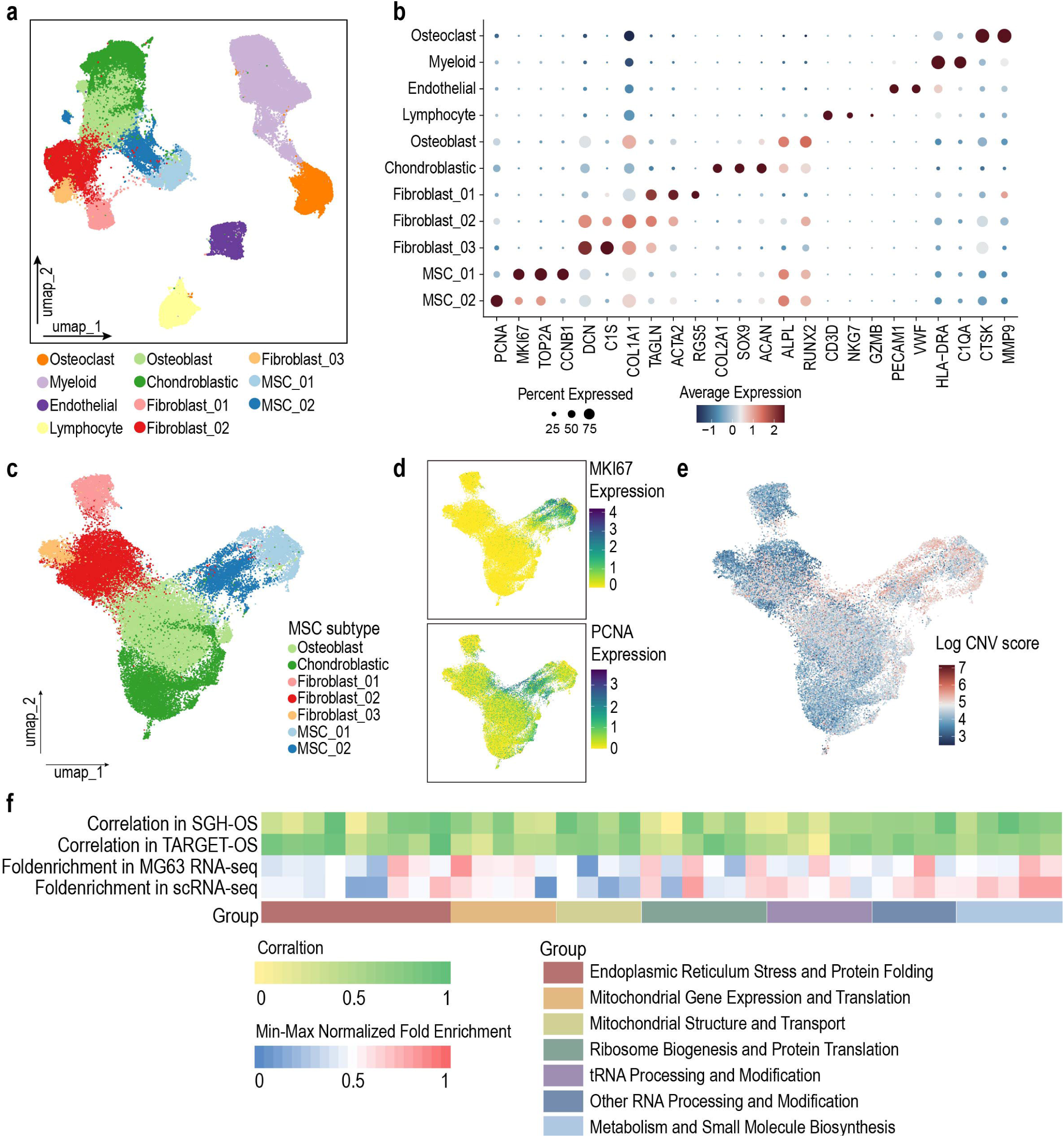

**Supplementary Figure 11.**
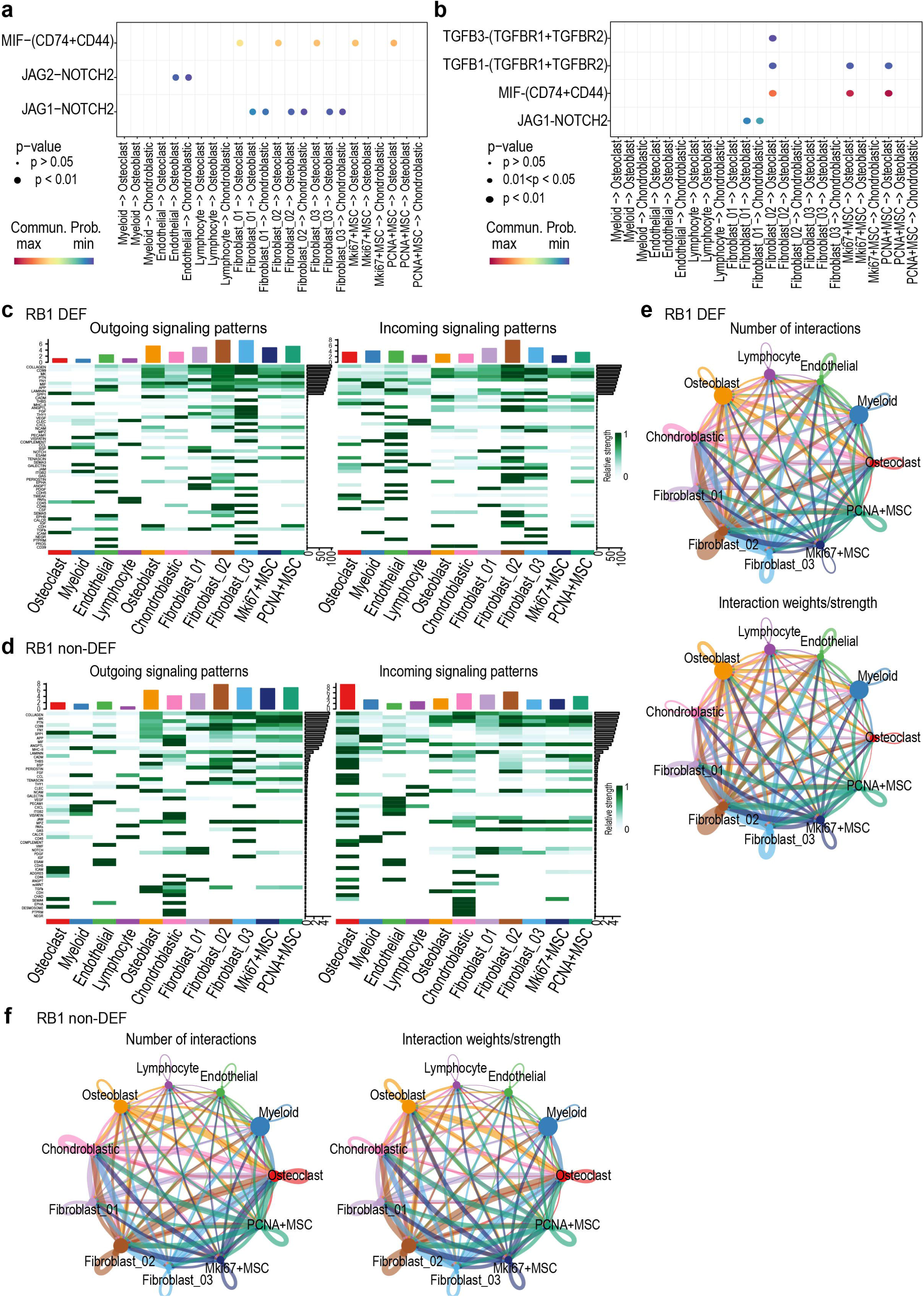

**Supplementary Figure 12.**
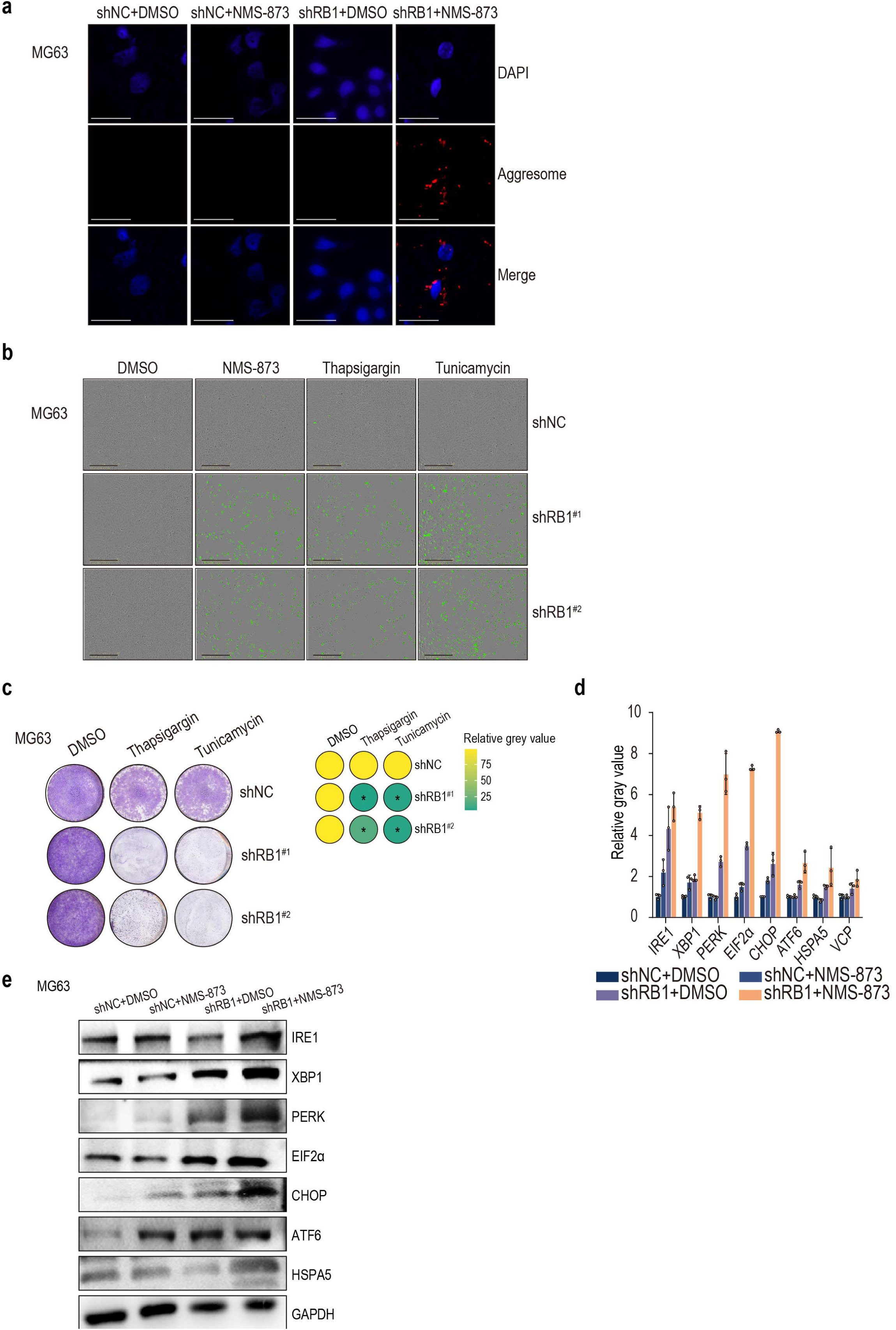

